# The Academic Career Readiness Assessment: Clarifying training expectations for future life sciences faculty

**DOI:** 10.1101/829200

**Authors:** Laurence Clement, Jennie B. Dorman, Richard McGee

## Abstract

We describe here the development and validation of the Academic Career Readiness Assessment (ACRA) rubric, an instrument that was designed to provide more equity in mentoring, transparency in hiring, and accountability in training of aspiring faculty in the life sciences. We report here the results of interviews with faculty at 20 U.S. institutions which resulted in the identification of 14 qualifications and levels of achievement required for obtaining a faculty position at three groups of institutions: research-intensive (R), teaching-only (T), and research and teaching-focused (RT). T institutions hire candidates on teaching experience and pedagogical practices, and on their ability to serve diverse student populations. RT institutions hire faculty on both research and teaching-related qualifications, as well as on the ability to support students in the laboratory. R institutions hire candidates mainly on their research achievements and potential, which may limit the diversification of the life science academic pathway.

## Introduction

### The mentor role of research faculty and potential barriers to success for aspiring faculty

In the life sciences, the success of aspiring faculty, graduate and postdoctoral (GP) trainees, is highly reliant on the scientific training and the professional development provided by faculty at research-intensive institutions. In particular, GP training often relies primarily on the *ability* and *knowledge* of one faculty member at each training level (graduate and postdoctoral) to serve as a mentor for aspiring faculty. Traditionally, mentors are expected to provide psychological and emotional support to the mentee, support the mentee in setting goals and choosing a career path, transmit academic subject knowledge and/or skills, serve as a role model (1). In addition, strong references from graduate advisors and postdoctoral Principal Investigators (PIs) are essential for faculty candidates to attain positions. As a result, these research faculty are responsible for providing aspiring faculty with 4 essential resources: 1) the information needed to identify the skills they should prioritize to attain their career goals; 2) the learning environment and opportunities to acquire these skills; 3) the assessment of the trainee’s progress, and feedback for improvement; 4) the letters of recommendation (and ideally, the sponsorship) for faculty positions.

The reliance of GP training on the *ability* of faculty to mentor trainees towards these faculty positions is of particular concern because there is evidence of faculty mentorship bias towards less represented populations of trainees (2, 3). In fact, mentoring is one of the main barriers reported by scientists from underrepresented groups transitioning from postdoctoral training to faculty positions at R1 universities in science, according to a National Institutes of Health survey (3). Several studies have also shown gender bias in letters of recommendations and candidate evaluation in academic hiring (2, 4, 5). In addition, as most research faculty have only experienced the research-intensive (R1) faculty career path, they may have limited *knowledge* of the skills required to attain other types of faculty positions (for example, faculty positions at Liberal Arts Colleges or Community Colleges), a challenge when supporting trainees with diverse academic career goals. In this sense, the GP training system has left out a component of diversity: diversity of faculty career goals. As a result, the importance of trainees’ reliance on mentors can create systemic inequities that could be especially detrimental to students of diverse demographics and career goals.

### Leveling the playing field: making faculty hiring criteria transparent and accessible to all trainees

To level the playing field among trainees, we aimed to develop an assessment tool that could allow trainees and mentors to assess trainee career readiness for diverse faculty careers regardless of the trainee or the mentor’s prior knowledge and pedagogical expertise. This instrument would measure the academic career readiness of trainees, providing a chance for them to receive formative feedback on their progress toward career-based training goals. As a validated instrument, the tool could also provide faculty hiring committees with a way to standardize their hiring processes.

We chose to develop the instrument as a rubric instead of a set of Likert-type items, as rubrics provide many advantages. Specifically, the rubric we sought to design aimed at supplementing some of the resources provided by mentors, as follows:

1. ***To provide transparency to aspiring faculty around faculty candidate evaluation criteria*** (resource 1, above). Rubrics are structured to delineate clear evaluation criteria, a key recommendation in inclusive education which, if expanded to graduate academic career preparation, could level the playing field between trainees with different levels of support from their mentors, and different prior knowledge (6–8).
2. ***To allow trainees to identify and prioritize training opportunities that will help them reach their career goals*** (e.g. the type of teaching experience needed, the type of funding opportunity they should apply for). Along the same lines, trainees could use a rubric to assess the potential of a particular learning environment (a laboratory when choosing a thesis laboratory, or an institution when searching for a postdoctoral laboratory) in providing them with the training opportunities they will need to reach their career goal, resulting in a better “match” for their goals (resource 2).
3. ***To provide trainees with the structure to receive formative feedback***, an essential feature of inclusive education (resource 3). The rubric can be used to structure discussion sessions with a mentor to identify skill gaps and develop a training plan tailored to the trainee’s faculty career of choice. This can be especially helpful for research faculty mentoring trainees targeting non-R1 faculty positions. In the absence of such discussions, the trainee can also use the rubric as a self-assessment tool and to inform an individual development plan (6, 9).
4. ***To standardize the evaluation process of faculty candidates.*** In education, rubrics are commonly used when multiple evaluators are involved (7, 10). For example, research mentors could use the rubric to structure their letter of recommendation for a faculty candidate and provide a specific and nuanced description of the faculty candidate’s abilities (resource 4). Having multiple research mentors base their recommendations on a common rubric could improve the standardization of the hiring process for faculty hiring committees who may wish to mitigate known biases of letter writers (4, 5). Lastly, hiring committees could use a rubric to structure their evaluation of candidates and address their own biases during the hiring process, including during hiring deliberations (11).
5. ***Evaluation of training programs:*** A final type of evaluation purpose for an academic career readiness rubric would be to help funding agencies and institutions evaluate the effectiveness of interventions aimed at improving the preparedness of trainees for faculty positions (12).

### Using evidence-based practices to develop an academic career readiness assessment rubric

To develop our instrument, we adapted instrument development methods described in the literature and used by others (13–16). However, this study stands out methodologically from classical instrument development studies because the literature defining our construct, *academic career readiness,* is limited, and definitions vary widely across fields. In the educational measurement field, *career readiness* refers to the readiness of high school graduates entering job training (17). In the human resources field, *work readiness* of graduates is defined as the “extent to which [college] graduates are perceived to possess the attitudes and attributes that make them prepared or ready for success in the work environment” (18). To our knowledge, there is no relevant body of literature that addresses academic career readiness, particularly in the context of aspiring faculty in the life sciences. Therefore, in addition to developing and validating an instrument, this study focused on identifying the attributes and characteristics that can be used to define and operationalize the life science academic career readiness construct by asking the following research questions (13): *How is academic career readiness defined? What qualifications and levels of achievements are required of faculty candidates?*

This study provides important findings on the qualifications required for obtaining a tenure-track faculty position in the life sciences at a wide range of U.S. institutions, as well as on the levels of achievement necessary for each required qualification. It also provides a blueprint for developing career readiness rubrics across career types and disciplines. An added value of the resulting ACRA rubric is that it can be used by funders and administrators to assess outcomes of training programs and by hiring faculty to standardize the faculty hiring process.

## Materials and Methods

### 1. Instrument Design

The overall instrument design methodology used to develop and validate the Academic Career Readiness Assessment (ACRA) rubric is presented in Table 1. It follows multiple stages of instrument validation which involve 1) reviewing the literature and consulting with experts to begin to define and operationalize the construct, 2) conducting expert interviews to establish internal validity of the instrument, and 3) pilot testing the rubric to continue collecting evidence of validity related to the relationship with external variables (13–16). Our study focuses mainly on the first two stages of instrument development and begins to address the third stage.

**Table 1:** Stages of development and validation of the ACRA instrument, and versions of the ACRA rubric produced at each stage (13)

| Stage | Objective | Research Question | Data Sources | Product |
| --- | --- | --- | --- | --- |
| 1. Substantive stage | Defining and measuring the construct. | *How is academic career readiness defined? What qualifications and levels of achievements are required of faculty candidates?* | Multiple sources, feedback from career and professional development experts. | Prototype ACRA |
| 2. Structural stage | Establishing content validity & response process validity. | *Does the rubric measure what it intends to measure? Does it accurately represent how faculty evaluate the academic career preparedness of candidates?* | Qualitative and cognitive interviews with life science faculty who had participated in faculty hiring in their department. | Final ACRA |
| 3. External stage | Establishing relationships with other variables (institution type). | *Do different types of institutions define academic career readiness differently? Do they require different qualifications and levels of achievement from faculty candidates?* | Pilot testing of Final ACRA rubric in a survey format with faculty sample involved in stage 2. | Final ACRA with hiring levels |

The first stage of validation consisted of defining the academic career readiness construct in the life science field (Table 1). This stage led to the development of a “prototype” rubric with ten qualifications, or *evaluation criteria*, and four levels of qualifications, or *quality levels* (6, 19, 20). The second stage of validation “*assists in the refinement of the theoretical domain*” (13). It involved interviewing experts, life science faculty who had participated in faculty hiring in their department, to assess the content validity of the items, as well as conducting response process validity through verbal probing (15). Findings informed the modification of the prototype rubric and resulted in the final ACRA rubric. The third stage of validation consisted of making explicit “*the meaningfulness or importance of the construct”* through *“a description of how it is related to other variables*” (13). Here, the external variables referred to the different categories of institutions in the U.S., with a focus on three groups of institutions that emerged from the first stage of validation: research-focused, teaching-focused and research- and teaching-focused institutions.

In fact, an important difference between this study and classical instrument development studies is the fact that there appears to be significant differences in hiring practices across institutions that *belong to the same category.* As a result, classical rubric validation methods, which are mostly based on Interrater Reliability (IRR) present some limitations. Therefore, in order to gain a deeper understanding of the differences between hiring patterns among different institutions, we used a combination of qualitative and quantitative data to begin assessing the external validity of the rubric, and found important differences between hiring patterns among different institutions.

### 2. Development of the Prototype ACRA rubric

In 2014, we began by developing a “Prototype” ACRA rubric using multiple sources, described in Supplementary Materials S1. These sources were specifically selected to meet the needs of our trainee population: biomedical graduate students and postdoctoral scholars aspiring to obtain faculty positions. Because a large majority of our student population are biomedical life scientists, we selected sources related to cell biology, microbiology, developmental biology, neurobiology, biochemistry, genetics, molecular biology, systems biology, immunology, stem cell biology, physiology, as well as general biology.

We began by reviewing job search advice books to locate potential evidence of hiring practices that would go beyond the advice by one or two faculty members. We also consulted the peer-reviewed literature around developmental frameworks and hiring practices related to the life sciences (8, 21–23). Although this step provided information on the types of qualifications required and some of the possible levels of achievement expected, the limited evidence-based literature relating to life science hiring practices across institutions prompted us to conduct a review of job postings.

In the Fall 2014, we collected and reviewed job postings that represented hiring practices at R1, Master’s granting universities and liberal arts colleges from online websites used by our trainees for applying to faculty positions. Here again, we focused on the specific subfields that were relevant to our trainee population, cited previously. We reviewed three R1 job descriptions, one Master’s (M) granting institution job description, three Liberal Arts College job descriptions (LAC) and found that the information provided in these job listings only provided limited information on the qualifications required, but alone were too limited to develop detailed levels of achievement for all these qualifications.

Our next step was to informally ask faculty in our network for feedback on the current applicant’s application materials. Three trainees who were applying to faculty positions volunteered their application packet. A convenience sample of three faculty (two R1 and one Liberal Arts College) in our network were emailed and invited to provide anonymous feedback to these three candidates via email (two faculty) and via Skype (one). These faculty belonged to the subfields mentioned earlier, and two had extensive experience hiring faculty at their institution. The third faculty member was a pre-tenured faculty who had a more recent experience of the hiring process, having been on the interview circuit, in addition to their experience reviewing applications for new hires at their new institution.

In addition, we leveraged the extensive expertise of career and professional development experts in our Office to inform the development of the rubric. This was done informally throughout the Fall 2014 application season by listening in to one-on-one CV review counseling appointments between a senior career advisor and biomedical trainees who were embarking on the faculty job market. In addition, the developer of the ACRA rubric, L.C. had experience as a faculty at a community college and used this experience to represent the hiring practices of community college faculty.

Later, when the first draft of the Prototype ACRA rubric was developed, it was presented informally to career and professional development experts in a group meeting and in one-on-one discussions to gather feedback on possible modifications to the framework. To ensure it was useful and understandable by trainees, it was also presented to graduate students and postdoctoral scholars in a professional development workshop as part of a module around the requirements of faculty positions. A survey was sent to trainees as a follow-up to determine the usefulness of the rubric in its current format. The resulting Prototype ACRA is described in the results section.

### 3. Development of the Final ACRA rubric

IRB approval: This project was done in compliance with the University of California, San Francisco Institutional Review Board Study # 15-17193.

#### Sampling

An initial sample of life sciences faculty members was selected using a criterion sampling strategy, based on the faculty members’ known interest in mentoring students and supporting diversity in higher education (24). From this initial sample, 22 faculty members with experience hiring tenure-track life science faculty were selected based on their institution’s Carnegie classification, using a maximum variation sampling strategy: to ensure that this rubric was inclusive of all tenure-track faculty career goals, we sampled institutions across research-intensive institutions, comprehensive universities, liberal arts colleges, and community colleges. The institutions were categorized according to the 2015 Carnegie Classification of Higher Education Institutions (25) as indicated in Supplementary Materials S2. Together, these Carnegie Categories represent 56.5% of all U.S. institutions and serve 88.3% of college students (25). The faculty represent diverse gender and ethnic/racial backgrounds.

Five of the faculty members were asked to participate in the pilot study, and 17 were asked to participate in the main study. Snowball sampling was used to identify an eighteenth faculty member. Together, these 18 faculty members had hiring experience at 20 different institutions.

#### Pilot faculty interviews

The pilot study was designed to 1) test and refine the verbal probes used in the interview guide and 2) determine if any essential qualifications had been omitted from the original rubric. Because our career and professional development experts were more familiar with large R1 and community college hiring practices, we selected five faculty who had experience hiring at primarily undergraduate institutions ranging from R2 to BAC, and a faculty who had observed hiring at a smaller R1 institution as a faculty. To collect meaningful feedback on the interview guide, we specifically selected faculty who had some familiarity with science education research design. In the first part of the interview, faculty were asked to answer the interview questions. In the second part, we asked faculty to describe their reaction to the interview process and to provide suggestions for improving the interview protocol. The pilot phase was conducted by L.C. and J.D. and resulted in the modification of the ACRA rubric to reflect feedback from faculty in the pilot group, and the modification of the interview guide, the development of an “Interview” ACRA rubric and an interview slide deck (Supplementary Materials S3).

#### Faculty Interviews

We then conducted semi-structured interviews with 18 additional life sciences faculty members who had experience hiring life science tenure-track faculty at 20 institutions. The interview guide involved asking faculty to select the factors that contributed significantly to hiring decisions at their own institution from among 16 qualifications. For each qualification they selected, faculty were then presented with four possible levels of achievement. We used verbal probing to assess the representativeness, clarity, relevance, and distribution of each selected qualification (content validity) and to assess comprehension of the items (response process validity) (15). In addition, the interview format allowed us to ask subjects to describe how they defined academic career readiness for this qualification in their own words, by describing the process for evaluating the qualification and describing the differences between candidates who were considered ready or not ready for a position at their institution. These data were later used to modify the rubric qualifications and level descriptions, as well as to begin to identify minimum hiring levels for each institution. Interviews were conducted by J.D. via video-conferencing, recorded, and audio was transcribed verbatim. One of the community college faculty was excluded from the analysis due to a malfunction in the recording process.

#### Analysis of faculty interviews and synthesis

The interview data was analyzed through a multistage process by adapting standard instrument development methods described in the literature and used by others (13–15, 26–30). Transcripts were first anonymized and coded by J.D., L.C, and a third researcher, using holistic codes based on the ACRA qualifications (31). Holistic coding consists of identifying sections of text and organizing them into broad topics “*as a preliminary step before more detailed analysis*” (31). To ensure that the final instrument accurately reflected the hiring practices of all types of institutions in our sample, we conducted a first round of analysis of the interviews by a group of institutions. Our initial hypothesis, based on the results of stage 1, was that research-focused, teaching-focused and research- and teaching-focused institutions prioritized different types of qualifications and required different levels of achievement for many qualifications. Therefore, our first cycle of analysis consisted of analyzing data separately for three groups of institutions: teaching-only institutions (T: Associate’s Colleges according to the 2015 Carnegie Classification, Supplementary Materials S2), research-intensive institutions (R: R1 Institutions), and research- and teaching-focused institutions (RT: R2, R3, M1, M2, Baccalaureate Institutions). In this first analysis cycle, sections of the interviews were analyzed across each institution group (R, RT, T) by researchers (J.D. and L.C.) and themes that emerged in each group analysis were used to modify the Interview rubric and create R, RT, and T “Group” ACRA rubrics. Group rubrics reflected the modifications described by faculty members as well as intra-group divergence in hiring practices. Reflective memos were developed for each group. To establish sufficient reliability of the analysis, J.D. and L.C. both reviewed all transcripts and discussed the new Group rubrics, including modifications and new qualifications identified by the other researcher to reach an agreement over the design of the final three Group rubrics. Next, we re-read all the transcripts, using the themes that had emerged in the group analysis. Analytical memos were developed to describe inter-group convergence and divergence. Based on these findings, the Group rubrics were synthesized to create one unique rubric (Final ACRA rubric, Table 5) that represented convergence and divergence in hiring practices over 4 levels. In addition, extensive footnotes were developed to help provide more details for users, a recommended practice in rubric development (6).

#### Testing with trainees

To determine if the rubric was clearly understandable by GP trainees, we used ACRA in multiple academic career development workshops with trainees who were at three career stages: 1) Faculty career exploration stage, 2) Skill-building stage (skills related to faculty positions), and 3) Faculty job search and application stage. In all these workshops, participants were UCSF biomedical graduate students and postdoctoral scholars, a large majority of which belonged to the life sciences subfields described in section 1 of methodology. Each time, trainees were presented with a short lecture summarizing the ACRA, given an opportunity to read the ACRA individually, pair, and share their questions. We also conducted one informal focus group with trainee volunteers during the development of the Final ACRA. In these multiple settings, participants were asked to share their questions about the rubric verbally and in writing. These questions were used to make minor changes to the language describing the levels, and improve the clarity of the accompanying footnotes.

### 4. Relationship with institutional type

The last stage of validation of the ACRA rubric involved 1) confirming that the Final ACRA rubric, developed as a result of the interviews, reflected the hiring practices of faculty (content validity), and 2) determining if it allowed to discriminate between the hiring practices of the three groups of institutions (external validity).

#### Sampling

The 23 faculty interviewed in the pilot study and in the main study were contacted via email and asked to complete a survey. Of these, 19 faculty responded. Two community college faculty, as well as two R1 faculty from the most elite institutions in the sample, did not respond to the survey. The community college faculty who had been excluded from the qualitative analysis was included here.

#### Survey design

Survey questions were identical to the interview questions but used the Final ACRA instead of the Interview ACRA. In the survey, respondents were asked to: “*Select from the list below the qualifications that contribute significantly to hiring decisions at your institution. Select only the qualifications without which a candidate could not be offered a faculty position at your institution*.”

The survey was developed using Qualtrics and was scaffolded to ensure higher participation rates. First, after selecting the essential qualifications from a list, respondents were presented with the description of the five levels solely for the qualifications they had selected and were asked to identify the *minimum achievement level required for candidates to receive a job offer in their department*. At the end of that section, they were given the option to see the rest of the qualifications (i.e, those that they had *not* selected from the list) and asked to identify the minimum level at which they hired, including Level 0 (is not a significant contributor to hiring decisions at this institution). In a third section, if they had selected “Prestige,” “Network,” or “Prior finding” from the list, they were also asked if they were willing to help define the levels for these qualifications (as these qualifications had not been selected by any faculty in the interviews).

To further establish response process validity and determine whether the new levels of achievement reflected actual hiring practices, we included a sixth response option for each qualification: “unable to assess” which triggered the display of an open-ended question prompting respondents to explain why they were unable to self-assess (13, 15).

#### Sample characteristics

Demographic data were collected at the beginning of the survey. 21% of faculty self-identified as ethnically or racially under-represented minority (URM) according to the National Institute of Health definition (American Indians or Alaska Natives, Blacks or African Americans, Hispanics or Latinos, Native Hawaiians or Other Pacific Islanders): 16% African-American and 5% Mixed race Latinx (32). In addition, 26% of respondents felt they belonged to groups underrepresented in research, beyond race and ethnicity: 16% as first generation to college, 5% as female and 5% did not provide any explanation. The sample included 42% male and 58% female faculty. Sixty-eight percent of faculty belonged to a biology department. The rest of the faculty belonged to departments closely related to biology (for example biochemistry), or to larger science departments that included biology faculty, or did not respond to the question. All faculty had observed at least 3 faculty hiring cycles in their department, with some faculty having participated in as many as 25 (Mean: 9.1, SD: 5.8, one “I don’t know” response). All but two faculty had served on hiring committees at their institution, with a range of experience from 2 to 17 hiring cycles (Mean: 6.3, SD: 5.0). Of these two faculty, one participated in the pilot study and one participated in the main study, and both were excluded from the survey data analysis. Contrarily to the qualitative study, each faculty were asked to only represent one institution’s hiring practices in their responses to the survey.

#### Analysis

Survey data was analyzed using Google Spreadsheets. To more clearly outline institutional differences and similarities, institutions were organized in groups. However, in the interest of making these findings useful for trainees, we used the new 2016 Carnegie Classification to categorize institutions (Supplementary Materials S4). With this new organization, all institutions belonged to the same group (R, RT, T) as originally described except for one institution, which moved from the RT category to the R category.

##### a. Reliability

Interrater reliability (IRR) is commonly used in instrument development to establish construct validity. Our goal was to use IRR as a way to determine the level of convergence between hiring priorities of faculty members at similar types of institutions (13, 16). We calculated IRR as percentage agreement between faculty from similar institutions (i.e. percentage agreement of faculty from R institutions was calculated, which was separate from the calculation for T faculty) (14, 16). When it came to the RT group, we assessed values for RT pairs of respondents from the same sub-type of institution (for example, we compared Baccalaureate colleges to each other). To ensure that we only compared data for faculty who had seen the entire ACRA scale for each qualification, we excluded respondents who had refused to complete the second section of our survey. This left us with 4 pairs of faculty, listed in Supplementary Materials S5. Because the scale had been designed to discriminate for intra-group differences in hiring practices, including intra-sub-group differences, we focused on calculating IRR agreement on whether the qualification was required to obtain a faculty position at that institution, not the minimal level of qualification of the institution. Divergent qualifications (qualifications for which faculty had divergences in agreement) are reported for each pair in *Results*.

##### b. Inter- and intra-group differences in required qualifications and minimal required level

Survey data were analyzed in two steps:

- **Step 1**: **“Required” qualifications.** For each group of institution, we calculated the percentage of institutions which selected each qualification as a significant contributor to hiring decisions, in two possible ways. In the first case, the faculty member had selected the qualification from the list of required qualifications, and when presented with the description of the levels, they did *not* select “level 0” (does not contribute to hiring decisions) in the menu of options. In the second case, the faculty member had not selected the qualification from the list but agreed to see the rest of the qualifications in the second part of the survey. Sixty-seven percent of T faculty, 63.6% of RT faculty and 67% of R faculty opted to see the rest of the survey, i.e. to assess the qualifications they had not originally selected from the list, and we included their responses to this second part of the survey here. Faculty members who did *not* select “level 0” for that qualification were included in the percentage calculations.
- **Step 2**: **Minimal hiring levels.** For each qualification, and for each group, we calculated the percentage of faculty who selected each hiring level as the minimum hiring level in the survey.

## Results

### 1. Development of the Prototype ACRA rubric

The first stage of this project involved the development of a Prototype ACRA rubric (Supplementary Materials S6), which represented our hypothesis for the operationalization of the academic career readiness construct. In the first step, we gathered and aggregated multiple sources to produce a draft of the rubric, which included nine qualifications (Supplementary Materials S7). In the second step, we requested feedback from career and professional development experts on the draft rubric to establish content validity of the instrument. The feedback received from these experts resulted in the addition of a tenth qualification, “Diversity Outreach” (Supplementary Materials S8). To create the four levels of achievement for this new qualification, we reviewed the resources described in Supplementary Materials S1 further. We also made some slight modifications to the format of the rubric as a result of the feedback, and the names of the qualifications to add specificity. For example, “Vision” became “Scientific Vision” and “Leadership” became “Scientific Leadership.” One of the major findings of this stage was that definitions of academic career readiness appeared to vary widely across institutions. To facilitate the presentation of the information to trainees, we categorized institutions into three main groups: research-intensive institutions, research- and teaching-focused institutions, and teaching-only institutions.

### 2. Development of the Interview ACRA Rubric

The pilot interviews led to the refinement of the interview guide and the modification of the Prototype ACRA. The resulting Interview ACRA rubric included 16 qualifications, including six new qualifications: Fit For The Position, Research Feasibility, Research With Undergraduates, Collegiality, Commitment To Diversity, and Personal Connections (Supplementary Materials S9). In addition, Teaching was redefined as two different qualifications: Teaching Experience and Teaching Philosophy, and the Leadership qualification was renamed “Independence.”

### 3. Development of the Final ACRA Rubric

#### 3. a. Defining academic career readiness: Which qualifications matter, and are there differences between institutions?

Faculty who participated in our qualitative study were presented with the list of 16 qualifications extracted from the Interview ACRA and asked to select those that contributed significantly to hiring decisions at their institution. These responses are summarized below and confirm our hypothesis that the three groups of institutions (R: research-intensive institutions; RT: research- and teaching-focused institutions; T: teaching-only institutions) defined in our study present distinct hiring profiles.

##### R hiring priorities

Our findings show that R institutions hire exclusively based on demonstrated research accomplishments and research potential of candidates. R faculty in our sample selected the same five core qualifications from the list: scientific vision, scientific independence, fundability, scholarship, and fit. Although some of the faculty selected scientific communication and research feasibility from the list, further probing into their importance in the decision-making process indicated that these two qualifications were not necessarily essential for a candidate to receive an offer for a position. In addition, R faculty explicitly said that the candidates’ mentoring and teaching skills, as well as their commitment to diversity, did not have any importance in the final hiring decision in their department: “*Yeah, so commitment to diversity: no. Teaching philosophy: no. Mentoring: no. Research with undergraduates: no.*”

##### RT hiring priorities

RT institutions compose a second, somewhat heterogeneous group with a distinct hiring pattern from R institutions. Faculty in this category selected collegiality, teaching experience and teaching philosophy, mentoring and research with undergraduates, and scientific communication as significant contributors to their hiring decisions. Some of the essential qualifications for R institutions were not relevant for this category of institutions, but research feasibility, fit, and scholarship were important.

##### T hiring priorities

The qualifications selected from the list by T institutions were: teaching experience, teaching philosophy, commitment to diversity, fit and collegiality. The cluster of qualifications selected by T institutions overlapped somewhat with RT institutions, but not with R institutions, except for the “fit” qualification, which spanned all three groups.

#### 3. b. Levels of achievement: Definitions and inter- and intra-group divergence

After selecting the significant contributors to hiring decisions from the list of 16 qualifications, faculty were shown four levels of achievement for each of the selected qualifications. They were asked to identify the level at which they hire, and in particular to focus on the minimal level of achievement required for a candidate to be selected for each step of the hiring process and finally receive a faculty job offer. Faculty were also asked to discuss the language used in the description of the achievement levels, and whether they reflected their definition of this qualification. The findings of these interviews and the rationale for modifying the ACRA items are described below by qualification. When describing intra-group divergences below, we occasionally refer to the institution’s Basic category in the 2015 Carnegie Classification of Higher Education Institutions (25).

##### 1. Expanding “vision” to include strategy

At R and some RT institutions, candidates are not only evaluated on their ability to develop a short-term and long-term research plan, as originally hypothesized. At a minimum, hiring faculty are also asking for candidates to present an exciting research question, with a plan that demonstrates that the candidate has a clear sense of direction. In addition, the candidate must be able to describe explicit, feasible steps to achieve their goals in the first few years. In addition, at R institutions, the research question must be broad enough to provide direction for the next 5 to 10 years, but it must also fill an important gap in the field. However, to “make the cut,” candidates for R positions must demonstrate that their strategy involves smaller projects that are well thought through methodologically (Table 2A). Ideally, in addition, the candidate’s productivity record will demonstrate that they have the ability to lead this project independently.

**Table 2:** Affiliation of the faculty member and Illustrative quotes for each of the seven Final ACRA qualifications identified by R faculty as being significant contributors to hiring decisions in the qualitative study.

| Inst. | Illustrative quote |
| --- | --- |
| **Research vision and strategy** |  |
| R1-1 | A. *"That's what we're looking for: "What's the question?" Tell me what the most important questions in your field are and how your research plan is perfectly suited to answer them. Especially in the chalk talk, I would like to see somebody start with that. That's the person who's going to impress me... And you have to show me that you're going to be able to take those big picture questions and break them down into, 'If I do this experiment, it will answer that question.'"* |
| **Publications** |  |
| R1-3 | B. *"We don't discuss impact factors. We try very hard not to pay attention to the journals, but inevitably someone will say something about Science, Cell or Nature. But there will be other people in the committee [that] are like, "Dammit. That's not ... We're talking about the science," like: "Is this science important?" There's a lot of awareness, I think, about trying to really think about "What's the important science? How are they going to advance science in the future?" The expectation is that it's a high level, it's creative, it's important, it's potentially new. Yeah. Impact is all those things in that, I think, that context..."* |
| **Funding plan** |  |
| R2-2 | C. Interviewer: *"Is what you're saying basically that the funding track record, looking back, is less relevant maybe than the funding potential, if you imagine seeing this person's package come through a study section?"*<br>Faculty: *"I think that's 100% true because not everybody has the opportunity (...) to write grants. Right?"* |
| BAC-2 | D. *"Let's imagine somebody's in a huge lab where nobody writes any grants. Do we really penalize them for not having written grants? No, but if they write well, if the papers are well-written, and they're first author, that's a really good sign that they can write and collect their thoughts."* |
| M1-2 | E. *"If we see an applicant that has [a K99], chances are we don't actually look at them that seriously, because there's no way they would come here once they see... Number 1, if they have that kind of grant, with the teaching load, there's no way they're going to be able to do that grant. So, at some level, having that doesn't really help you here. It probably hurts you, but again, it's position-dependent."* |
| R3-1 | F. *"Fundability. Yep. That is another essential thing because we do hope that you secure outside funds and in the application, at least you've demonstrated that you've thought about how you might pursue these funds. Agency: have you looked into the funding by the agency, and so forth. Will they fund an undergrad institution? (...) At least they've shown that they've at least begun to research how their projects might be funded, like what agencies. (...) What's most likely or most appropriate sources of funding, (...) The more specific they are, the better; if they don't just say NSF or NIH, but they know particular programs- (...) The better. They seem more serious."* |
| R1-2 | G. *"Someone who has thought at least through the first major grant. Not, "I'm going to come in and this is the first experiment I'm going to do." It's: "This is this key important question that I'm going to spend the next 5 to 10 years of my life trying to understand." And a lot of problems that postdocs run into is they think of the next experiment...or set of experiments, and not, "The long-term goal of my research is ..." That should be a statement in your R01. And that needs to be big and important ...you would have to convince someone that it's worth doing."* |
| **Research independence** |  |
| R1-2 | H. *"Are they actually really dependent on people who do certain techniques or certain collaborators at their home institution, and will they be able to maintain those connections when they establish their own lab?"* |
| R1-1 | I. *"It's about: they were the ones who were thinking about what the next experiment would be. They were the ones who are developing the collaborations. They were the ones who were doing all that stuff."* |
| R1-3 | J. *"I would say that I think this idea of being independent, there is often a lot of discussion out of "is this person distinct from their postdoc advisor?" If they seem similar, is the advisor letting go of a project or was it a new direction within?"* |
| R1-3 | K. *"It ranges from, "We've talked about this and all of these things are going to mine to take," could be, "I'm really good friends with my advisor and we communicate regularly, and I know we're going to do that in the future. We're not going to step on each other's toes." That's what I said in my case, I was like "yeah, we communicate, we have communicated, we'll keep communicating." People need to have thought about ... If they're like, "I don't know," that's a bad sign."* |
| **Recommendations** |  |
| M1-1 | L. *"I think there's a big red flag if the person doesn't get letters from certain people that they should be getting letters from (...) If they don't have that, they really should explain it. (...) Because that might make us feel better. Otherwise we're just going to ignore them (...) maybe it would work as a side e-mail to the chair of the committee. (...) Because most of the time people just don't say anything, and we're left with: "this is kind of worrisome"."* |
| **Fit** |  |
| CC-1 | M. *"Like somebody who has an understanding and meets the needs of the institution would be somebody who has this understanding of equity and has teaching experience and has a teaching philosophy that matches those things. In a way this seems like a compound category."* |
| M1-2 | N. *"A lot of it's just fit for the position. Anyone that's too similar to what we already have, we typically remove, because we don't have the capacity for redundancy here. (...) I think disciplinary fit for us is probably the most important thing. That's where ... We have so few faculty to cover all of biology. We still have holes. We can't have redundancy."* |
| R2-1 | O. *"It's really like survival in a really challenging environment where you're going to have to be resourceful, you're going to have to be able to get along with a lot of different personalities, and you're going to have to launch a research program while also teaching at a very high level, and that you need to have a certain grit to do that, right?"* |
| R1-3 | P. *"Fit, I think for us, will sometimes mean within the context of our department or community ... Will this person provide something new scientifically? Will they synergize with different communities that exist? (...) some schools will try and build strength in an area, so I think fit can mean those things, as R1 institutions."* |

##### 2. Publications: reflecting the debate in the evaluation of impact

The “publications” item was modified to reflect the wide range of hiring levels found in our sample. We found that publications were not relevant in T hiring practices and that some of the more teaching-focused RT institutions, like some baccalaureate colleges, only required the production of a few papers, regardless of authorship or impact. Some RT institutions focused on the regularity of the candidate’s publication rate as a first author, often preferring regularity over impact. The major modification for this section of the rubric focused on the last two levels, and were designed to reflect a debate among the faculty interviewed over the evaluation of publication impact, and whether it should be assessed through the impact factor of the journal or whether the impact of the work on the field should be evaluated independently (Table 2B).

##### 3. Fundability: shifting the emphasis from prior achievements to the funding plan

Although most R faculty picked “Fundability” out of the list of qualifications, they indicated that prior funding was not how they assessed fundability. Faculty explained that they felt that a candidate’s funding record could depend on factors that do not solely reflect ability. For example, faculty believed that some trainees may not have the opportunity to write their own grants (Table 2C, D). Even in the case of the K99 funding mechanism, which is open to foreign nationals, there was some debate at both R and RT institutions on its value when selecting candidates (33). One R faculty recognized that K99 recipients may be hired preferentially by some institutions, but expressed reservations when it comes to this particular funding mechanism and its ability to prepare scientists for the R01 grant process. It is worth noting also that non-R faculty may disqualify candidates who do have a K99 grant (Table 2E).

When it comes to fundability, what seemed to matter for most institutions was the candidate’s plan for obtaining external funding. The importance of such a plan varied among institutions, and seemed to perhaps be based on the tenure requirements of the department.

Fundability was not an essential qualification at RTs, especially at institutions with minimal tenure requirements relating to extramural funding. In contrast, at RT institutions where obtaining tenure requires some extramural funding, the candidate is not only expected to propose a research program that can be funded, but also to demonstrate a basic knowledge of funding mechanisms for which they would be eligible (Table 2F). Finally, in RT departments where internal collaborations are frequent and where mentoring between faculty is expected, faculty will evaluate the fundability of a candidate on their *potential to work well with the colleagues who will help them get funded*.

The last three levels of the rubric reflect the important emphasis that R faculty put on the candidate’s funding plan. At these institutions, where extramural funding requirements for tenure are high, a research program needs to be ambitious enough to be eligible for R01 funding, and impactful enough to be appealing for reviewers in an R01 study section (Table 2G). This assessment of one’s ability to appeal to an R01 study section is done indirectly, by assessing the candidate’s research program through the lens of a funding agency, and/or explicitly, by asking the candidate to outline the aims of their first R01 grant. Faculty find that many candidates have not prepared adequately for this assessment, and either do not have any aim, or the aims are unrealistic, both deal breakers for a hiring committee. Candidates are also more attractive if they are able to outline aims for more than one large grant (in their chalk talk, for example).

##### 4. Leadership vs. independence: distinguishing oneself from one’s mentor matters, team management skills do not make a difference

Although R institutions selected “Scientific Independence” as a significant contributor to hiring decisions, their definition differed significantly from ours. Our original hypothesis was that scientific independence involved managing a research project and a research team independently. The R faculty members in our sample felt that managing a project independently was a skill they expected of the selected candidates, without explicitly selecting for it, except in situations where the work presented had been a large collaborative project. The type of research independence that R institutions are, in fact, looking for is mostly assessed at the last stage of the hiring process, at the chalk talk and during one-on-one interviews, although faculty members are searching for signs of independence as early as in the CV. Research independence, for R faculty, means three things:

1. ***Technical independence***: Does the candidate have the technical expertise to run their proposed research program independently? If the program relies on collaborations, will these collaborations be maintained in the new position (Table 2H)?
2. ***Independence of thinking:*** Did the candidates develop their own ideas independently? Because many of the candidates who make it to the last stage of selection come from large laboratories managed by prestigious PIs, there is a concern that the candidate may be simply implementing their PI’s vision (Table 2I).
3. ***Distinct “niche” from their PI:*** This aspect is directly related to fundability. If the candidate cannot explain how they will navigate the potential competition with their PI’s research program, they run the risk of competing for grant proposals, in which case it would be assumed that they would be less likely to succeed (Table 2J). A lack of clear answer to a question about the potential competition with the candidate’s PI is an important red flag when hiring faculty (Table 2K).

Although it isn’t necessary, ideally, faculty would like to see mentors and PIs reinforce the candidate’s independent vision and achievements in their letters of recommendation, which was added as a fourth level of the Research Independence qualification.

##### 5. Recognition vs. recommendation: specific and detailed is better, stellar is often required

We found in this study that letters of recommendation were determinant in getting trainees from the application stage to the in-person interview stage, but not as relevant in the final decision to make an offer. We also found that the levels we had originally developed did not represent the different hiring levels of institutions. At both R and some RT institutions, it would be a red flag if applicants came in without a letter of recommendation from their current, postdoctoral PI (Table 2L). Overall, search committees are looking for recommendations saying that the candidate has the attributes to become a successful PI, as defined in other sections of this study. For example, they look for recommendation letters that confirm the candidate’s independence from their PI’s research program. In addition, some Rs will require letters that affirm that the candidate shows the potential to become a leader in the field.

##### 6. Fit: addressing a spectrum of definitions

Interestingly, fit was selected as a significant contributor to hiring decisions by most faculty in our sample but we found that they had various definitions of fit. In fact, some faculty suggested that it may be a compound category (Table 2M).

- ***Institutional fit:*** Our results confirm our hypothesis that at some institutions, fit may mean demonstrating an understanding with the institutional mission. This understanding is best demonstrated by the candidate having sought out experiences that demonstrate alignment with the institution’s teaching and/or research mission (Table 2M). Candidates at RT institutions, in particular, will be expected to carefully tailor their application materials to the institution.
- ***Disciplinary fit:*** In addition to institutional fit, we found that, for many RT institutions, fit also means disciplinary fit, whether in research or in teaching. Because some of these institutions have limited capacity to hire faculty, they want to avoid disciplinary redundancies to ensure that all their classes are taught by faculty with experience in the discipline, a selection that happens early on in the application process. Avoiding redundancy of research programs is also a concern for R institutions (Table 2N).
- ***Professional fit:*** For some institutions, fit means having the resilience, the “grit” to work in a highly demanding environment with high teaching expectations and possibly significant research expectations (Table 2O).
- ***Potential fit and synergies:*** For R institutions, fit can also mean developing something new or synergistic with existing research or teaching communities at the institution (Table 2P).

##### 7. Verbal communication of research

All R faculty and four RT faculty selected “Scientific Communication” as a significant contributor to hiring decisions from the qualifications list. Faculty members at RT and at some RT institutions (R2, M1, see Supplementary Materials S2) required candidates to present their science clearly to faculty from different life sciences sub-disciplines than their own (for example, for microbiology candidates, presenting their science to ecologists). Although this was the minimum level of achievement required by R faculty (Table 3A), one faculty member did mention that a candidate who could only present their research clearly to scientists from their own subfield (i.e. microbiologists) could still receive an offer for a position in their department, suggesting a difference between hiring *intentions* and *actions*.

**Table 3:** Affiliation of the faculty member and Illustrative quotes for each of the Final ACRA qualifications identified only by RT faculty as being significant contributors to hiring decisions in the qualitative study.

| Inst. | Illustrative quote |
| --- | --- |
| **Verbal communication of research** |  |
| R2-1 | A. *"(...) there was always a fair amount of like the ecologists would have a voice and they'd have something at the table. They would be at the table in the discussions and so they needed to understand what was going on. If they said, "I have no idea," it was a concern."* |
| M2-1 | B. *"What we're mainly looking at is being able to explain it to undergraduates. (...) I think part of it is in an organization, that helps scaffold the information for the listener, so they're not jumping right to these acronyms and these concepts that some people start out that way, and they start out even in their summary, they feel like they have to get all the big words in there. (...) just being aware of the audience and maybe explain things more than you would think is necessary."* |
| **Research feasibility with available resources** |  |
| M1-2 | C. *"The vision, feasibility have to really be tied together there. They might have a great vision, but if they can't do it here, they're not going to be hired."* |
| M1-1 | D. *"Research feasibility definitely involves resource constraints(...) We have some really odd, amazing resources, so I think it's always in the best interest of the applicant to do their research. (...) I think [they would] have to email the chair because we have a really bad website. I'm sure that that could be potentially true for other places. You wouldn't want to depend on the internet."* |
| R2-1 | E. *"What I had in mind was more the institutional resource constraints. It was more like extremely expensive boutiquey instruments that might end up being like a five million dollar piece of equipment or something. (...) Then, concern about having that thing run and be serviced, and all of the sort of just support community that might need. Yeah, organisms, it was like anything would work and really same with microscopes. They were very open to any kind of microscope that was needed. It was a struggle sometimes if somebody needed the super duper, duper fancy two photon yaddah, yaddah that was \$1.2 million, but usually they got that. It was more the super high end stuff."* |
| R1-1 | F. *"Sometimes people who come from HHMI labs, giant labs, then they might have the wrong idea about how life is going to be. (...) how many postdocs would you need to do that? No, you're not going to have 25 postdocs in your first year. Do not think this is your boss' lab."* |
| **Inclusion of undergraduates into the research plan** |  |
| BAC-2 | G. *"We get about 100 applicants. Right off the bat, about 50 of them can be tossed because they didn't read the ad, or they don't know [our institution]. They say they look forward to working with our graduate students and postdocs, so we toss those."* |
| M1-1 | H. *"We ask that question, we basically say: "how would you involve undergraduates in your research?" Then we'd look at what they say. That's definitely one of the phone questions. It has always been. We don't dock them if they've never done it before, but we listen to how they would do it, and if it sounds like they've really thought it through carefully than they're good." (...) We definitely hire people who have thought very thoroughly on how they would involve undergrads, and they were convincing. (...) It really is there thoughtfulness based in reality. (...) Do they kind of realize that undergrads come and go?"* |
| R3-1 | I. *"Some of this comes up in the phone interview, one of the questions may be along the lines of, you have your research program, how would you define a Master's [student] project versus an undergrad research project?"* |
| **Experience conducting research with students** |  |
| R2-2 | J. *"To some extent, people care that you're interested in what the undergrad is trying to get out of the experience, and you're trying to meet that need. (...) I meet with my students weekly and we make these goals for the next week. We don't need to hear everything, but I like people to recognize that not one style works for every person. You might need to change your mentoring style."* |

In addition, the candidate’s ability to explain the science to undergraduates was essential for RT faculty and was added as a new level (Table 3B). In addition, some RT faculty indicated that candidates were not asked to offer a research talk but a *teaching demonstration* that does not focus on the candidate’s research field. Because this practice amounted to evaluating teaching practices and potential, we did not include it in the “Verbal Communication” qualification.

##### 8. Research feasibility with available resources

Like “Fit”, the “Research Feasibility” qualification was added to ACRA as a result of the pilot interviews, to reflect RT institutions’ limitations in terms of research resources (Table 3C). All RT faculty interviewed expected candidates to present a research program that was tailored to the resources available at their institution, which may require contacting someone at the institution to inform the research program proposal (Table 3D). At some RT institutions, this meant that the system used by the candidate needed to also be low-cost and easy to use. At one very research-focused RT and one slightly less prestigious R (as well as one R in our pilot sample, a smaller institution in size) resource constraints amounted to obtaining very costly high-end equipment or access to specific patient samples (Table 3E).

However, even at R institutions where resource constraints are not an issue, some candidates struggle with adjusting their vision to the resources of a new lab. For example, hiring faculty were particularly worried about candidates who trained in a well funded Howard Hughes Medical Institute (HHMI) lab, because they may not realize what was realistically feasible without HHMI funds (Table 3F). There were also concerns about the candidate lacking the technical ability to manage an expensive piece of equipment, if their postdoctoral lab had the funds to support staff to maintain such equipment.

##### 9. Inclusion of undergraduates into the research plan

Unlike at R institutions, at many RT institutions, the integration of students into the research program was one of the most important hiring criteria, and as a result, we added a new evaluation criterion in the rubric to reflect the candidate’s ability to include the needs of undergraduates into their research plans.

At the most basic levels, institutions reported wanting to see that candidates had a clear understanding that they would be working with non-PhD students (Table 3G). Candidates also need to understand the impact of the constraints of these students’ scheduling availability and laboratory competency levels (novices vs. advanced students) on the scope of the project: “*Who’s going to carry their cell lines if they have to study for an exam? It’s clear that you have to think about the constraints of undergraduates, and that they’re full-time students*,” explained M1-2. At the next level, candidates should show that they have thought carefully about how they could involve students in their research (Table 3H). Finally, at the highest level, candidates will be expected to suggest projects feasible by the different populations of students enrolled at the institution, from Master’s degree students to freshmen (Table 3I).

##### 10. From “mentoring” to “experience conducting research with students”

We found that, at the most basic level of achievement of the mentoring qualification, candidates must articulate a scientific mentoring philosophy that meets the needs of the student population served by a given institution (Table 3J). In addition, having experience conducting research with students is a plus. Interestingly, some RT faculty reported that, even when they did not select for mentoring qualifications, their candidate pool presented the qualifications.

The next level of achievement involves producing preliminary data while conducting research with students, followed by presenting these data in a presentation or published article: *“I would say the hiring minimum is that posters… that they need to have not just had undergrads, but they should have mentored them all the way through presentations.”* BAC-2 faculty.

##### 11. Teaching experience

Teaching experience was not a significant contributor to hiring decisions at any of the Rs or at one of the most research-focused RTs. In addition, two RT faculty (belonging to R3 and M1 institutions, see Supplemental Materials S4) reported that having too much teaching experience or demonstrating too much passion for teaching *could* work against the candidate if the scholarship wasn’t strong enough (Table 4A).

**Table 4:**
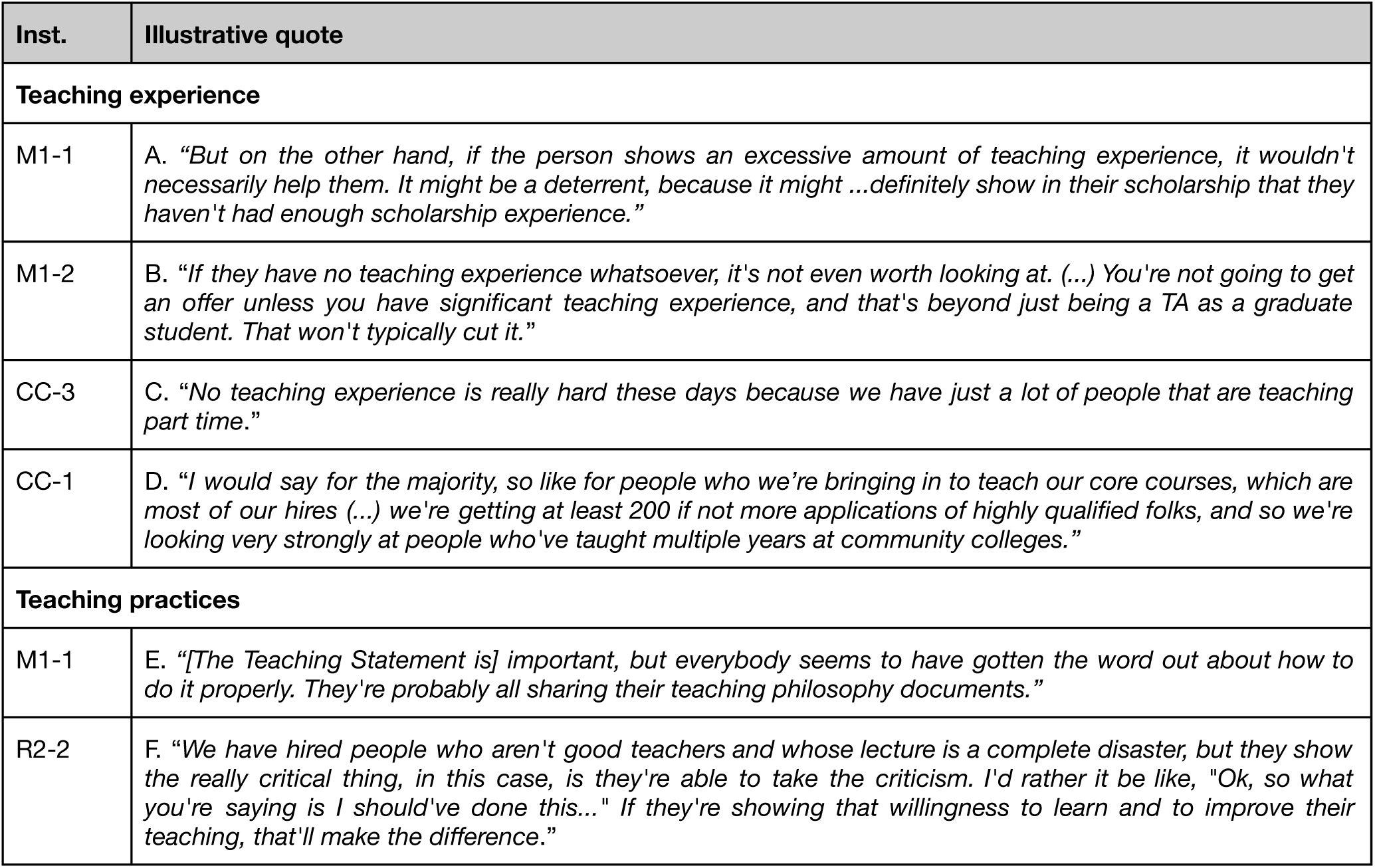
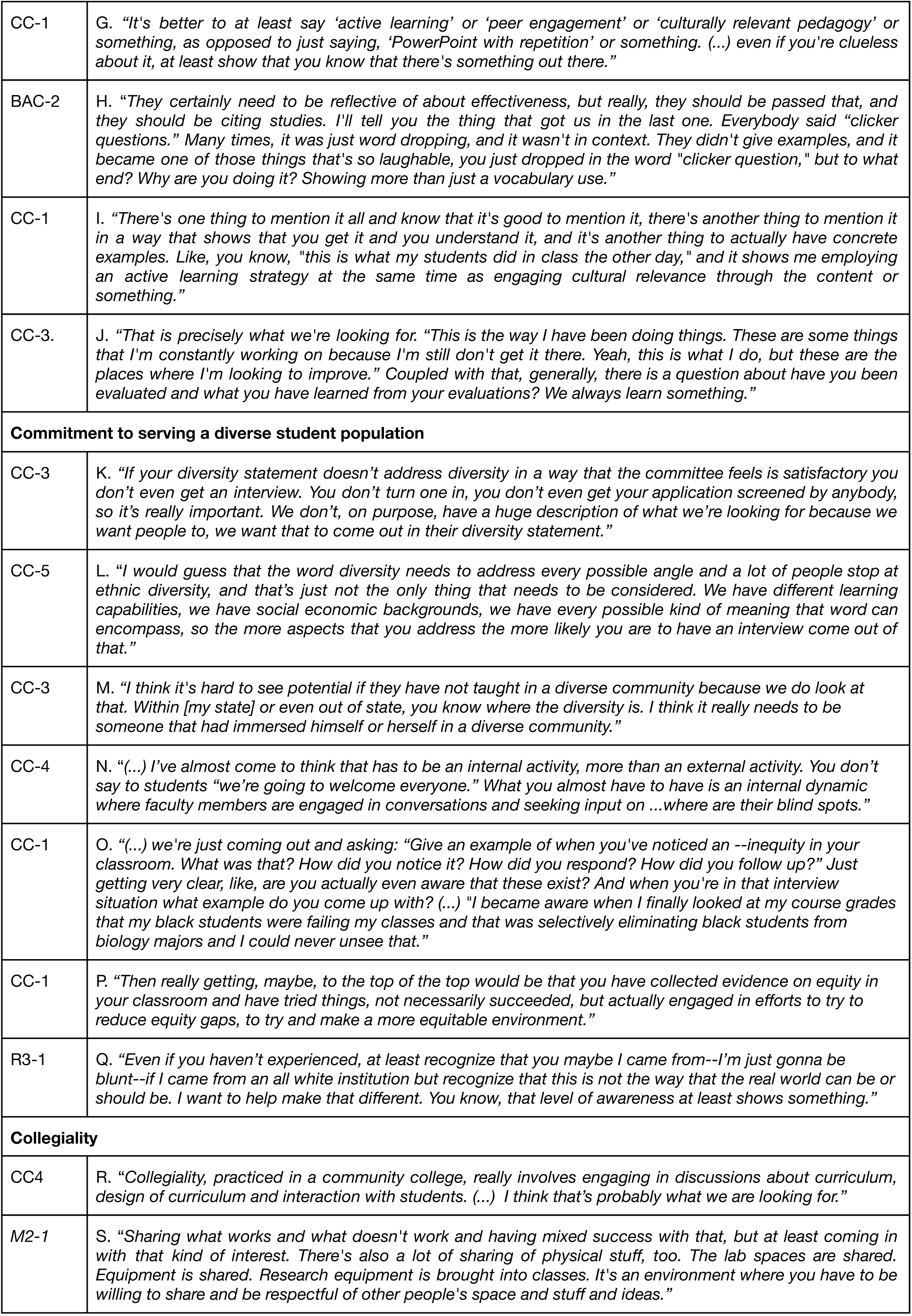
Affiliation of the faculty member and Illustrative quotes for each of the Final ACRA qualifications identified by RT and T faculty as being significant contributors to hiring decisions.

| Inst. | Illustrative quote |
| --- | --- |
| **Teaching experience** |  |
| M1-1 | A. *"But on the other hand, if the person shows an excessive amount of teaching experience, it wouldn't necessarily help them. It might be a deterrent, because it might ...definitely show in their scholarship that they haven't had enough scholarship experience."* |
| M1-2 | B. *"If they have no teaching experience whatsoever, it's not even worth looking at. (...) You're not going to get an offer unless you have significant teaching experience, and that's beyond just being a TA as a graduate student. That won't typically cut it."* |
| CC-3 | C. *"No teaching experience is really hard these days because we have just a lot of people that are teaching part time."* |
| CC-1 | D. *"I would say for the majority, so like for people who we're bringing in to teach our core courses, which are most of our hires (...) we're getting at least 200 if not more applications of highly qualified folks, and so we're looking very strongly at people who've taught multiple years at community colleges."* |
| **Teaching practices** |  |
| M1-1 | E. *"[The Teaching Statement is] important, but everybody seems to have gotten the word out about how to do it properly. They're probably all sharing their teaching philosophy documents."* |
| R2-2 | F. *"We have hired people who aren't good teachers and whose lecture is a complete disaster, but they show the really critical thing, in this case, is they're able to take the criticism. I'd rather it be like, "Ok, so what you're saying is I should've done this..." If they're showing that willingness to learn and to improve their teaching, that'll make the difference."* |
| CC-1 | G. *"It's better to at least say 'active learning' or 'peer engagement' or 'culturally relevant pedagogy' or something, as opposed to just saying, 'PowerPoint with repetition' or something. (...) even if you're clueless about it, at least show that you know that there's something out there."* |
| BAC-2 | H. *"They certainly need to be reflective of about effectiveness, but really, they should be passed that, and they should be citing studies. I'll tell you the thing that got us in the last one. Everybody said "clicker questions." Many times, it was just word dropping, and it wasn't in context. They didn't give examples, and it became one of those things that's so laughable, you just dropped in the word "clicker question," but to what end? Why are you doing it? Showing more than just a vocabulary use."* |
| CC-1 | I. *"There's one thing to mention it all and know that it's good to mention it, there's another thing to mention it in a way that shows that you get it and you understand it, and it's another thing to actually have concrete examples. Like, you know, "this is what my students did in class the other day," and it shows me employing an active learning strategy at the same time as engaging cultural relevance through the content or something."* |
| CC-3. | J. *"That is precisely what we're looking for. "This is the way I have been doing things. These are some things that I'm constantly working on because I'm still don't get it there. Yeah, this is what I do, but these are the places where I'm looking to improve." Coupled with that, generally, there is a question about have you been evaluated and what you have learned from your evaluations? We always learn something."* |
| **Commitment to serving a diverse student population** |  |
| CC-3 | K. *"If your diversity statement doesn't address diversity in a way that the committee feels is satisfactory you don't even get an interview. You don't turn one in, you don't even get your application screened by anybody, so it's really important. We don't, on purpose, have a huge description of what we're looking for because we want people to, we want that to come out in their diversity statement."* |
| CC-5 | L. *"I would guess that the word diversity needs to address every possible angle and a lot of people stop at ethnic diversity, and that's just not the only thing that needs to be considered. We have different learning capabilities, we have social economic backgrounds, we have every possible kind of meaning that word can encompass, so the more aspects that you address the more likely you are to have an interview come out of that."* |
| CC-3 | M. *"I think it's hard to see potential if they have not taught in a diverse community because we do look at that. Within [my state] or even out of state, you know where the diversity is. I think it really needs to be someone that had immersed himself or herself in a diverse community."* |
| CC-4 | N. *"(...) I've almost come to think that has to be an internal activity, more than an external activity. You don't say to students "we're going to welcome everyone." What you almost have to have is an internal dynamic where faculty members are engaged in conversations and seeking input on ...where are their blind spots."* |
| CC-1 | O. *"(...) we're just coming out and asking: "Give an example of when you've noticed an --inequity in your classroom. What was that? How did you notice it? How did you respond? How did you follow up?" Just getting very clear, like, are you actually even aware that these exist? And when you're in that interview situation what example do you come up with? (...) "I became aware when I finally looked at my course grades that my black students were failing my classes and that was selectively eliminating black students from biology majors and I could never unsee that."* |
| CC-1 | P. *"Then really getting, maybe, to the top of the top would be that you have collected evidence on equity in your classroom and have tried things, not necessarily succeeded, but actually engaged in efforts to try to reduce equity gaps, to try and make a more equitable environment."* |
| R3-1 | Q. *"Even if you haven't experienced, at least recognize that you maybe I came from--I'm just gonna be blunt--if I came from an all white institution but recognize that this is not the way that the real world can be or should be. I want to help make that different. You know, that level of awareness at least shows something."* |
| **Collegiality** |  |
| CC4 | R. "Collegiality, practiced in a community college, really involves engaging in discussions about curriculum, design of curriculum and interaction with students. (...) I think that's probably what we are looking for." |
| M2-1 | S. "Sharing what works and what doesn't work and having mixed success with that, but at least coming in with that kind of interest. There's also a lot of sharing of physical stuff, too. The lab spaces are shared. Equipment is shared. Research equipment is brought into classes. It's an environment where you have to be willing to share and be respectful of other people's space and stuff and ideas." |

However, for the rest of the RT and T faculty in our sample, candidates without any teaching experience were often disqualified because of the levels of experience of the candidate pool (Table 4B, C). In fact, guest lecturing and teaching assistantships, the type of teaching experiences usually provided at R institutions for graduate and postdoctoral scholars, were insufficient for candidates to be offered positions at many RT and T institutions (Table 4B).

It is noteworthy, however, that the level of specialization of the position may influence the size of the applicant pool, and therefore the amount of teaching experience that candidates need to be competitive for the position: “*I’ve been on a microbiology hiring committee where that’s a bit more specialized and we get a lot less applicants, and so there, we might be happy to have someone who’s just taught at university level, you know, microbiology or something, but that’s pretty rare.” - CC-1 faculty*.

Yet, faculty report that a growing number of candidates have several semesters of teaching experience, often with the type of student population the institution serves, an important advantage. For example, competitive RT candidates may have a year of experience as a Visiting Assistant Professor at another primarily undergraduate institution after their postdoctoral or graduate training. For T candidates, experience as a part-time Adjunct Faculty at a community college is often required as a demonstration of the candidate’s commitment and ability to teach community college students (Table 4D). Faculty in our sample recommended that R1 trainees interested in a faculty position at an RT or T institution should consider adjusting their training plan to gain teaching experience at an institution that serves the type of population they hope to serve someday.

##### 12. From teaching “philosophy” to “practices”

From our interviews, we found that some RT institutions struggle to find true value in the teaching philosophy statement, and use it primarily to vet applicants who may not be aware of the teaching culture at their institution, an evaluation criterion already described in “Fit” (Table 4E). In reality, many of the RT and T institutions assess teaching practices, or the potential to use certain practices, throughout the different stages of the interview and, when applicable, the teaching demonstration. At the most basic level, RT and T faculty are looking for some indication that candidates are aware that their teaching skills may not be as advanced as that of their colleagues, but that they are interested in developing these skills with some mentoring (Table 4F).

At the next level, faculty require at the minimum some awareness of the existence of evidence-based pedagogical approaches and, preferably, that teaching is a field that is being actively researched (Table 4G, H). In this case, candidates should be able to explain how student-centered approaches can be used effectively in the classroom (Table 4G, H). Some faculty would also hope to see candidates who can demonstrate that they can *use* these strategies. In fact, when the interviewing process involves teaching demonstrations, the candidate has an opportunity to demonstrate how they would use active learning in a classroom, even if they have little prior teaching experience (Table 4I). Ideally, candidates should also have experience teaching with student-centered approaches and the ability to reflect on successes and failures in teaching to adjust their teaching strategy and to improve their curriculum (Table 4J).

##### 13. Commitment and ability to serve a diverse student population

###### Struggles with defining the qualification

This qualification was selected by all T faculty members, except for one. Only two of the RT faculty (and none of the R1 faculty) selected this qualification as significant. Interestingly, the selection of candidates on their commitment to serve diverse student populations was not necessarily related to the diversity of the student population. For example, an RT faculty representing two BAC institutions serving a high proportion of underrepresented students reported that neither institution purposefully screened for the candidate’s commitment to diversity. Because faculty also misunderstood “commitment to diversity” as meaning that the institution was *committed to hiring faculty members from diverse backgrounds*, we renamed the qualification “Commitment and ability to serve a diverse student population.”

At T institutions, the candidate’s understanding of diversity issues is an essential part of the hiring decision, and the screening takes place through a thorough evaluation of the diversity statement at a very early stage of the hiring process (Table 4K). T faculty defined diversity beyond racial and ethnic diversity and included socioeconomic, cultural, education and career stage, career goals, first-generation status, and learning preferences, among other characteristics, and expected successful candidates to use broad definitions as well (Table 4L).

###### Evaluating abilities to serve diverse student populations

Although we had hypothesized that the candidate’s ability to teach and mentor diverse student populations would be a relevant skill, faculty reported that the teaching and mentoring experience in itself did not necessarily demonstrate that the candidate had the appropriate level of expertise. In reality, faculty at T institutions preferred to look for a candidate’s experience immersing themselves in a diverse community (Table 4M). As an illustration of this, T faculty reported that candidates with teaching experience at a community college had an advantage over other candidates. Specifically, T faculty were looking for candidates who had the basic sensitivity and respect for, as well as the interpersonal skills to interact with individuals of all backgrounds and felt that community college teaching experiences could demonstrate this skill. In addition, some faculty reported looking for a demonstration that the candidate had *used* teaching practices that help all types of students succeed in their classroom. For hiring faculty, supporting diversity in the classroom as an instructor involved being reflective about one’s own biases as an instructor or thinking about ways to adapt teaching strategies to the diverse needs of the students, instead of having students adapt to the instructor’s practices (Table 4N). An example of the strategies committees look for involves maintaining equity in the classroom by eliciting participation of all types of students. This use of strategies is evaluated at all stages of the hiring process, from the diversity statement to the teaching demonstration and may even be taken into account when administrators select a candidate from a list of recommendations by the committee (Table 4O). One of the community college faculty members also provided an additional level of evaluation, which involves expanding the “diversity” definition to social justice and equity, and emphasizes fairness for all students, one basic tenet of evidence-based teaching (Table 4P). More qualified candidates can describe a personal experience with social justice and equity in education or research, and in particular, experience with the marginalization of students that has impacted their teaching and mentoring practices. This experience can be direct (experienced) or indirect (observed). This level of achievement was required by only one T institution and was considered an advantage for other institutions (Table 4Q).

##### 14. Collegiality: the potential for working well with colleagues

Collegiality, in this study, was defined as the candidate’s ability to interact well with colleagues, and the potential to establish internal collaborations. In this qualitative wave, it was identified as a significant contributor to hiring decisions at both RT and T institutions, but it was not relevant to R institutions, mostly because such a competency was of significant importance in departments where resources are scarce, or where faculty need to work together to support non-traditional students.

At a minimum, candidates need to be able to interact with colleagues in a professional manner. At the next level, faculty described assessing candidates for their interpersonal skills to fit in the department culture. We kept the language of this achievement level broad enough to include the differences in institutional cultures. For example, at R institutions, the candidate would demonstrate curiosity for other faculty’s work and ideas, while at RT institutions, the candidate should be able to interact well with others, and get along with their colleagues. At T institutions, this fit with the departmental culture is an emergent category, that involves demonstrating skills in teaching and a commitment to serving diverse student populations (Table 4R).

RT and T institutions have a preference for candidates who are collaborative: who can work well with others, and who are willing to share ideas and resources with colleagues (Table 4S). Finally, candidates who demonstrate the potential for developing collaborative research and/or teaching projects with colleagues are more competitive for positions at T and RT institutions.

#### 3.c. The Final ACRA rubric

As described above, the analysis of the transcripts resulted in the modification of the list of ACRA qualifications and of the description of their corresponding levels, which are synthesized in the final version of the ACRA (Table 5). The resulting rubric comprises 14 qualifications with 5 levels that represent all levels of hiring by all institutions in our sample.

**Table 5:**
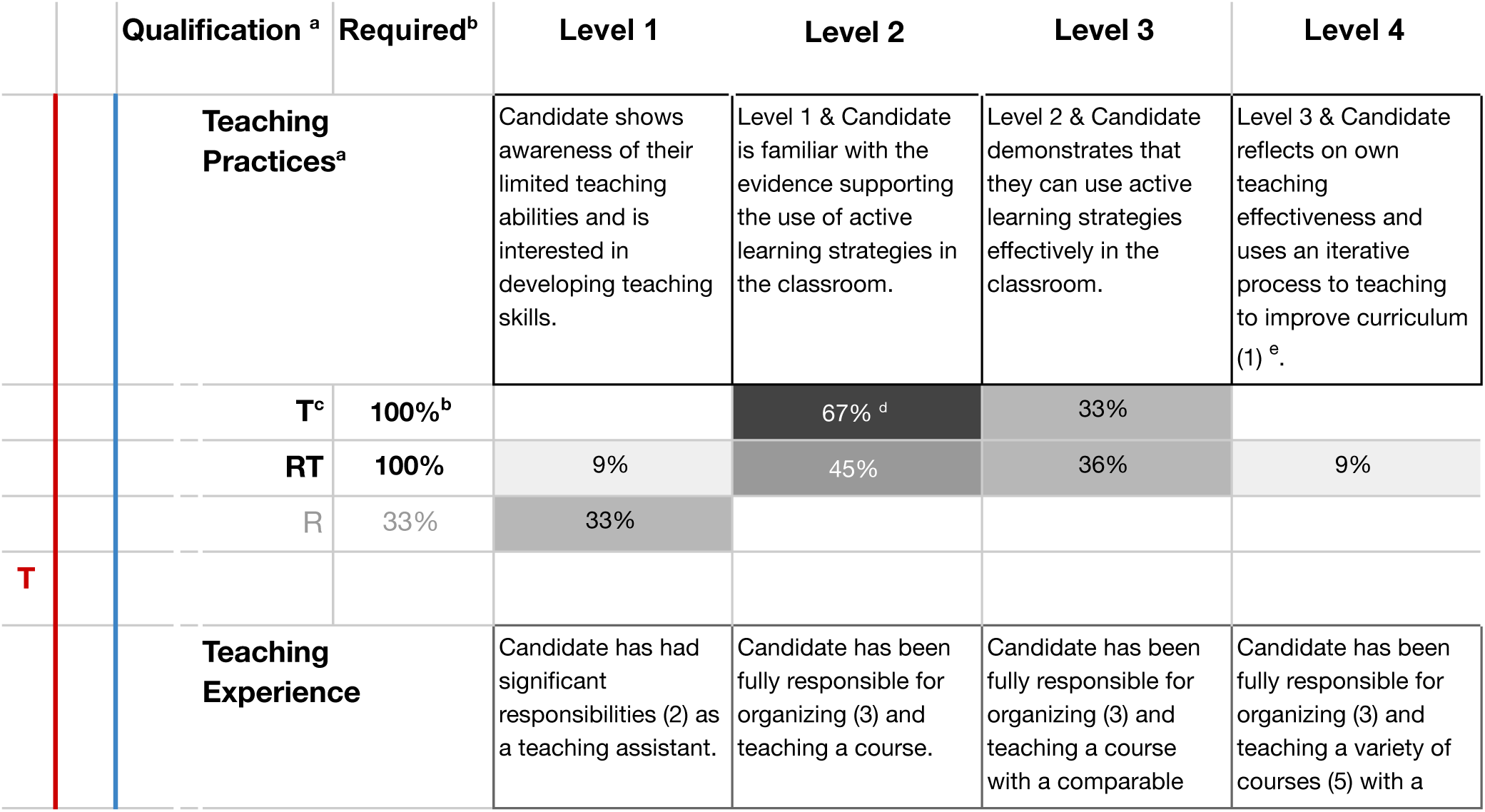
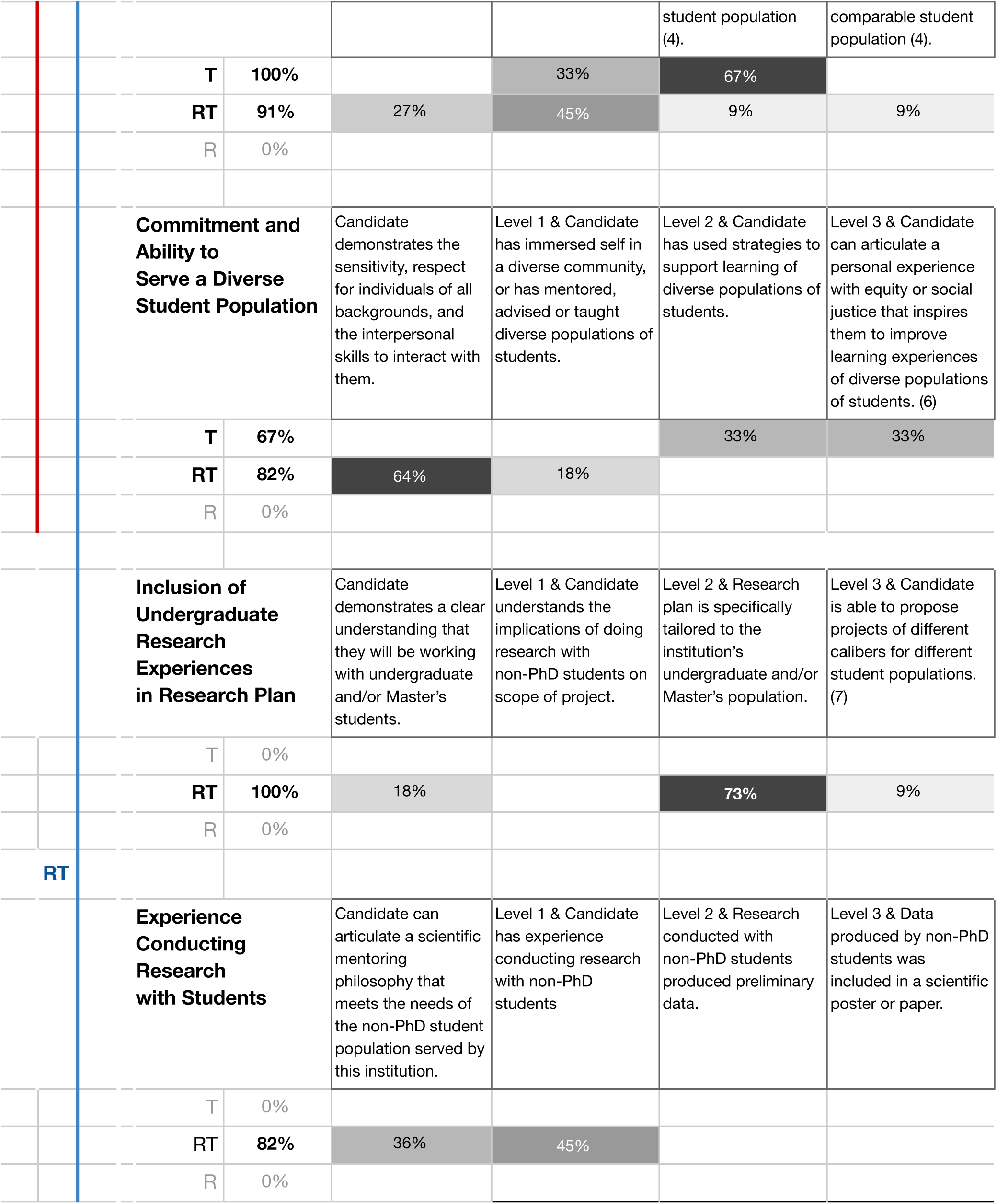
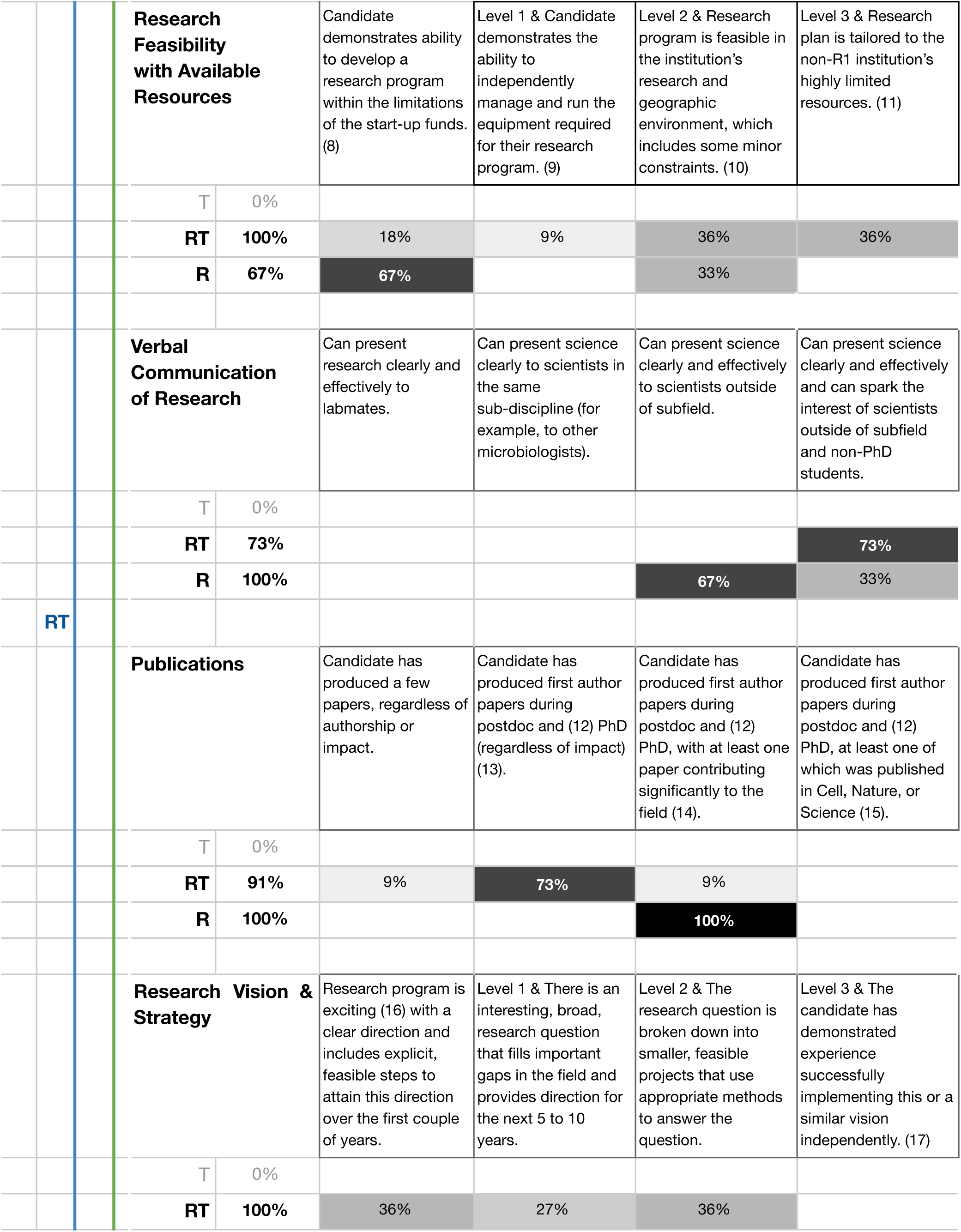
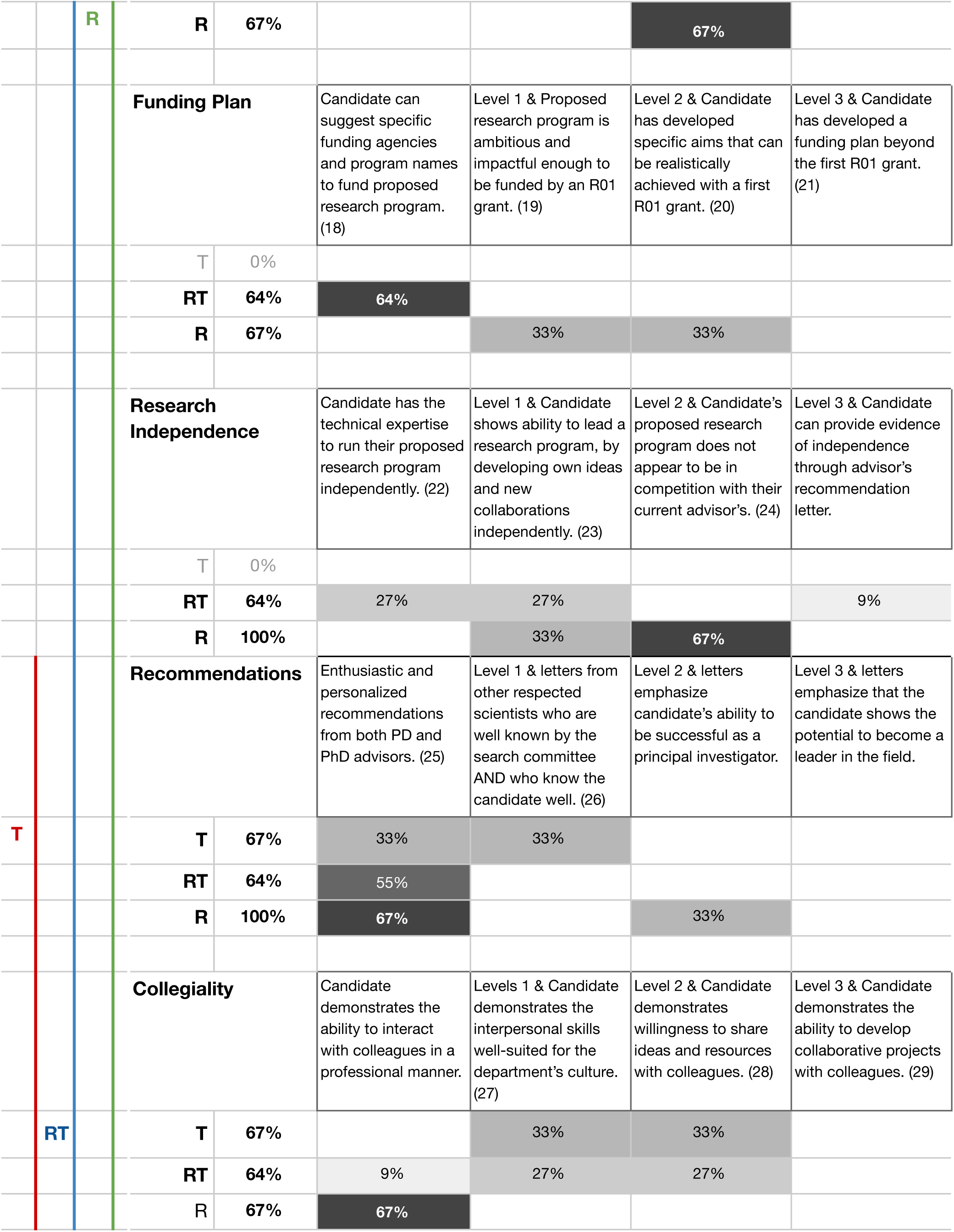
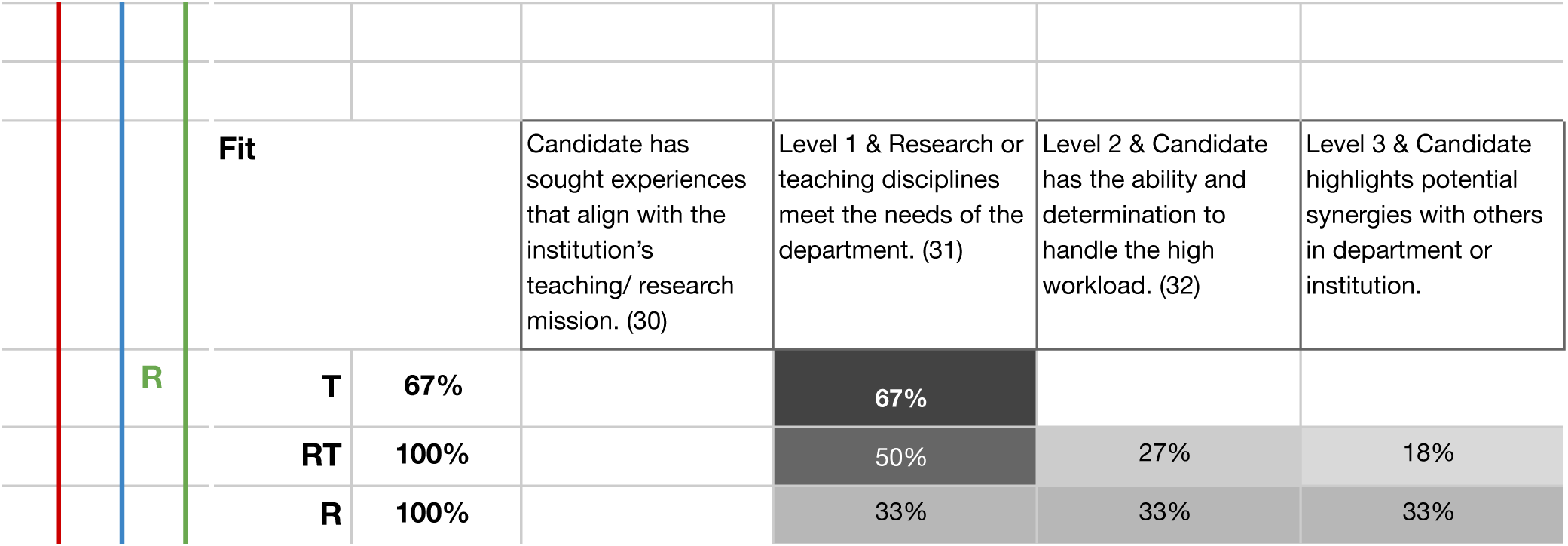
The Final Academic Career Readiness Assessment (ACRA), including the 14 qualifications that were selected as required by at least 50% of institutions in at least one group of institutions (R, RT, T). **^a^** “Qualification” refers to the rubric evaluation criterion, describing each of the qualifications identified in the qualitative study. Four of the five levels of achievement are represented here (Level 0 describes the absence of selection for each qualification). Faculty responses to the online survey using the Final ACRA rubric are represented under each qualification. **^b^** “Required” refers to the percentage of faculty from each institution group who selected the qualification as a significant contributor to hiring decisions. **^c^** Categories of institutions correspond to the “basic” classification in the 2016 Carnegie Classification of Higher Education Institutions (25): T (n=3); RT (n=11); R: (n=3). Bold font is used when at least 50% of faculty members in that institutional category selected the qualification. Color lines indicate required qualifications for each group of institution (T: red line, RT: blue line, R: green line) **^d^** Minimal hiring levels: Percentages in grey boxes represent the proportion of faculty who selected the rubric level as the minimum level required for a candidate to obtain a position at their institution. ^e^ Numbers in parentheses refer to footnotes in Supplementary Materials S10. The full ACRA rubric with accompanying descriptive footnotes can be downloaded from career.ucsf.edu/ACRA.

### 4. Differing hiring priorities and achievement levels required for R, RT and T institutions

In the third part of our study, we piloted the Final ACRA rubric in a survey format to determine if a) the rubric effectively reflected the hiring requirements of faculty in our sample, b) the rubric could detect intra-group similarities, and c) the rubric could reflect the inter-group and intra-group differences in required qualifications and minimal required achievement levels.

#### 4.a. Does the Final ACRA rubric reflect definitions of academic career readiness across institutions?

To ensure that the Final ACRA rubric reflected all qualifications and levels of achievements required at all types of institutions in our sample, we tested the following rival hypotheses.

##### 1) Rival Hypothesis 1: Some of the Interview ACRA qualifications omitted from the Final ACRA are, in fact, significant contributors to hiring decisions at one of the three groups of institutions

Survey respondents were given the option to select qualifications that they had *not* selected in the qualitative study from the list of qualifications. To that effect, four qualifications not included in the Final ACRA but included in the Interview ACRA were added back into the survey: a) Collaborations, b) Network, Professional Connections, c) Pedigree (Reputation of Training and/or Education), and d) Prior fellowships and grants. One RT faculty selected “Collaborations” from the list, but when provided with the scale, indicated it was not a contributor to hiring decisions. None of the respondents selected “Network,” three RT faculty members selected “Pedigree,” and one RT and one R faculty members selected “Prior fellowships.” When presented with the “Collaborations” scale in the second section of the survey, one T faculty selected Level 2, suggesting that collaborations were significant contributors to hiring decisions at their institution. Together, these findings show that only a minority of faculty members in one or two institutional groups identify these qualifications as requirements for faculty positions at their institution, demonstrating that the Final ACRA rubric is representative of the construct of academic career readiness.

##### 2) Rival Hypothesis 2: Some qualifications are missing from the Final ACRA rubric

Respondents had the option to select “other” from the list of qualifications. Two respondents selected “other,”. A T faculty member indicated “minimum degree qualifications,” and an RT faculty member indicated “willingness to teach undergraduates,” a qualification that is already included in the Final ACRA rubric. These findings support the hypothesis that the Final ACRA rubric qualifications accurately represent the academic career readiness construct.

##### 3) Rival Hypothesis 3: Final ACRA rubric level descriptions do not reflect all hiring practices of R, RT and T institutions in our sample

Respondents were provided with the option to indicate that they were unable to assess their required level of achievement based on the levels provided to them. None of the respondents included in our study selected this option. These findings demonstrate that the levels of achievement developed in the Final ACRA reflect all levels required by the institutions in our sample.

#### 4.b. Is the Final ACRA rubric a reliable instrument for assessing required qualifications at different types of institutions?

A common measure of validity used in rubric development is inter-rater reliability (IRR), where instruments with IRR values of 80% or higher are considered reliable (16). Our results show a high IRR score for the Final ACRA rubric in all three groups of institutions (T, RT, and R), from 80% for Baccalaureate colleges (RT group) to 100% for Master’s colleges (Larger programs, RT group). All qualifications had agreement between raters in at least two groups of institutions in the Final ACRA rubric (Supplementary Materials S5).

#### 4.c. Is the ACRA survey tool effective at detecting intra- and inter-group differences in hiring practices?

Table 5 presents the intra- and inter-group differences in academic career readiness definitions based on the survey data. The results confirm our qualitative findings in that T institutions select candidates principally on their teaching experience, teaching practices, commitment and ability to serve students from diverse backgrounds, collegiality and fit for the institution. In addition, we found here that recommendations may play a role in T faculty hiring as well. In this case, the T faculty is most likely referring to teaching-related recommendations: “*Someone that had been a teaching assistant and had just excellent recommendations.*” - CC-3

Our findings that RT practices overlap with T practices are also confirmed here, as well as the fact that RT institutions have lower requirements then T institutions when it comes to the level of skill expected from candidates in supporting diversity. In addition, we found that a majority of RT institutions also shared hiring practices with R institutions. Finally, we confirmed that RT institutions focus on three qualifications that are unique to their group, which involve student research experiences, and working with limited research resources. As with the T group, we found from these new data that recommendations mattered at RT institutions, albeit less than at R institutions.

When it comes to R institutions, our findings that R institutions hired principally solely on research-related qualifications were confirmed, with the novel finding that collegiality of the candidate mattered for two out of three of the R institutions surveyed.

These findings show that the Final ACRA rubric can detect differences in hiring patterns within each group of institutions, a first step in developing further studies of predictors of hiring decisions.

## Discussion

### 1. Significant contributors to hiring decisions reflected in the ACRA rubric

#### Teaching-Only Institutions (T)

Overall, T faculty in our sample had the highest requirements when it came to the candidate’s commitment and ability to serve diverse student populations, and stood out from the RT and R institutions in our sample when it came to the prioritization of these skills in their hiring decisions. At most T institutions, candidates must not only demonstrate the respect for individuals of all backgrounds and have mentored, advised or taught students from diverse backgrounds, they must also demonstrate that they have used strategies to support learning of diverse populations of students in class or in the lab. In some instances, candidates must also be able to articulate a personal experience with equity or social justice that inspires them to improve the learning environment of diverse populations of students. A majority of the T faculty in our sample also required a significant amount of teaching experience from candidates, who must demonstrate that they have been fully responsible for organizing and teaching a course with community college students. T faculty also expected candidates to not only be interested in improving their teaching skills, but also to be familiar with the evidence supporting the use of active learning strategies in the classroom and, in some cases, demonstrate that they can use these active learning strategies effectively in the classroom. An important selection criterion for T faculty was whether the candidate could teach the disciplines required by their department. Because T faculty are not expected to take on any research-focused responsibilities, research potential and accomplishments were not evaluated as part of the T faculty hiring process.

#### Research-Intensive Institutions (R)

R faculty in our sample selected candidates mainly on demonstrated research accomplishments and potential of candidates. Candidates must demonstrate a solid and regular first-author publication record, with highly impactful work in their field. Depending on the R institution, candidates may not be required to have published in the highest impact factor journals (Cell, Nature, or Science). Candidates are also expected to propose a research plan that is independent of their PI’s work, with a clearly delineated research vision and feasible strategy that is of the caliber of an R01 grant. Candidates must be able to communicate their research clearly to scientists outside their subfield, and, in some institutions, communication of science to non-PhD students is also required. Letters of recommendation from graduate and postdoctoral advisors are essential across Rs. In addition, at some institutions, candidates must have letters from scientists who are well known by at least some members of the hiring committee and know the candidate well, and these letters must emphasize candidate’s ability to be successful as a principal investigator. Disciplinary fit with the position is also a significant contributor to hiring decisions, and at some institutions, it may also be necessary for candidates to highlight potential synergies with others in the department or institution.

Although R faculty are almost always expected to take on teaching responsibilities, and are in charge of mentoring all future faculty and PhD-level scientists in the scientific workforce, candidates’ commitment to serving diverse student populations, their teaching experience, their teaching practices, and their undergraduate mentoring experience or potential do not appear to be significant contributors to hiring decisions.

#### Research and Teaching-Focused Institutions (RT)

A majority of the RT faculty in our sample required significant teaching experience, including having organized and delivered at least one course. Candidates are expected to demonstrate that they had held curricular responsibilities (syllabus, lecture, assignment and exam development) and had developed classroom management skills by organizing and teaching a course. At a subset of RT institutions, candidates would be required to demonstrate more teaching experience, similar to the level expected at T institutions. In addition, 90% of RT faculty in our sample required an understanding of evidence-based teaching practices and close to half of them required experience using these practices in an effective manner. Over 80% of RT faculty expected candidates to present a research plan that described how they included undergraduate or Master’s students in their research. Another way in which candidates needed to signal their interest in an RT position was to propose a research plan that was tailored to the limited start-up funds and institutional resources. Candidates must also be able to spark the interest of non-PhD students and scientists outside of subfield when presenting their research. A majority of the faculty in our sample required first-author publications during the graduate and postdoctoral training, regardless of the impact of these papers. The candidate’s research vision and strategy were important to a large majority of the RT institutions, with variable levels of achievement required, sometimes at the level of an R institution. Prior undergraduate mentoring experience was a requirement at a subset of RT institutions, but most required that candidates describe a mentoring philosophy that would meet the needs of their students. Disciplinary fit and the ability of candidates to work synergistically with colleagues, while handling the high workload was also evaluated by RT institutions.

### 2. Implications and future work

This work has many implications, with impact on a) the development of future faculty, b) the graduate career education research field and c) the diversification of the academic pipeline. In this section, we also describe ways in which the science education research and the research communities can come together to refine and expand ACRA beyond its current scope.

#### a. Impact on the development of future faculty

The ACRA rubric provides a higher level of transparency to the faculty hiring process for trainees in the life sciences. It can be used by trainees with limited mentor or institutional support to advance their careers. In addition, R1 mentors can use ACRA to better guide their mentees in developing the skills they need for the faculty position of their choice. Graduate career educators and faculty can use the ACRA rubric to inform trainees on the types of faculty positions available to them, develop an individual development plan, and help them showcase their relevant skills in faculty application materials.

Many graduate career educators around the country are already using the ACRA rubric with their trainees. At our institution, the ACRA rubric has been used in individual advising appointments to help trainees assess their preparedness for faculty positions and identify goals for their discussions with mentors. The Final ACRA rubric has been used to develop a list of recommended training goals for aspiring faculty, tailored to the type of institution targeted by trainees. We have used an evidence-based practice, backward-design, to strategically plan and develop our academic career development programs in alignment with these ACRA-based training goals (34, 35). To provide teaching experience to our GP trainees, we offer teaching residencies in partnership with local RT and T institutions, and we provide training in teaching practices through an evidence-based pedagogical course as well as a Science Education Journal Club, both of which also help candidates develop the skills to support diverse student populations in the classroom (36–38). To provide trainees with the skills to support diversity in the laboratory, we have developed a course focused on inclusive mentoring, supervising and educating practices for laboratory researchers as well as mentoring residencies with community college students (39, 40). Because of the importance of letters of recommendation in the faculty hiring process, our Office has also been developing a series of programs to help trainees navigate the relationship with their PI and other mentors (41).

In addition, we developed an ACRA-based workshop aimed at helping early-stage trainees make academic career decisions and develop a training plan based on these training outcomes (42). For late-stage trainees, an additional series of ACRA-based workshops, including application material samples and peer-review checklists, were designed to guide participants in their faculty application process (43). Life science trainees targeting RT institutions as well as non-life science trainees are encouraged to use the ACRA rubric to structure their informational interviews and better understand the hiring requirements of the types of positions which they would like to obtain. We are now in the process of developing a graduate-level course mapped to ACRA R1 training goals to help trainees attain less well-known qualifications required for R1 faculty positions, such as developing an independent research vision.

#### b. Impact on the field of graduate career education research and program evaluation

This study represents an effort to translate effective education research practices to the field of graduate career education, expanding the use of Discipline-Based Education Research (DBER) to the academic pipeline beyond undergraduates (44). To our knowledge, this study is the first of its kind to attempt to define in a systematic manner the construct of academic career readiness for future STEM faculty using a rigorous instrument validation method. Additionally, it provides a blueprint for the definition and operationalization of career readiness across disciplines, career types and education levels. The Graduate Career Consortium Education Research Committee is currently working on developing an ACRA rubric for humanities disciplines and will soon begin developing a rubric for non-academic careers for life scientists (a “NACRA”) (45).

This study also supports the growing trend for funding agencies to require solid evidence that trainees are receiving adequate support and are progressing toward pre-established training goals (46). This need for evidence has prompted graduate programs and grant-funded faculty development programs to search for solid assessment tools to evaluate their success. The ACRA rubric is one such tool that could be used to assess training programs or interventions in a systematic and longitudinal matter.

#### c. Impact on the diversification of the academic pipeline

Recent studies have found that to increase the proportion of underrepresented minority (URM) faculty, graduate and postdoctoral training programs should focus efforts on the *transition from trainee to faculty* (47, 48). To improve this transition, the life science education and research fields must begin to consider interventions that can address the career-development of future faculty: “*we must broaden the focus of professionalization and rectify the imbalance between training for research and training for a career”* (49, 50). In recent years, some interventions have been used to supplement the research faculty mentoring role, and multiple governmental funding mechanisms aimed at supporting the needs of trainees with diverse backgrounds and diverse career goals (51–55). In addition, many institutions have taken steps to provide additional support to trainees by creating positions that are focused on delivering supplemental instruction, as well as career and professional advising to trainees (45, 56).

Unfortunately, these resources are not accessible to all trainees. As an additional source of career information that does not solely rely on mentor knowledge and ability, ACRA presents the potential to improve equity in the development of future faculty across institutions and mentoring opportunities by diminishing the likelihood that their mentor is a trainee’s only source of career information. It provides clearer expectations for aspiring faculty, with the potential for each trainee to define their own training goals based on systematically collected evidence of hiring practices.

However, diversifying the academic pipeline also requires that the scientific community 1) address the biases in hiring practices and 2) equip faculty and mentors with the skills to mitigate their own biases in the classroom and in the laboratory.

If confirmed with a larger cohort, the findings that R1 faculty are hired exclusively on their research accomplishments potentially unearth some important barriers to the latter. If a faculty candidate’s mentoring experience, use of evidence-based teaching practices and commitment, or ability to serve diverse student populations are not significant contributors to hiring decisions at R1 institutions, the under-represented trainees whom they will serve as faculty will be at a disadvantage in the classroom and in the laboratory.

The result of ignoring these student-related qualifications in R candidates could create a vicious circle where faculty are unequipped to hire other faculty who are prepared to support institutional diversity. For example, we have noticed that R1 trainees and faculty are often confused with what diversity statements are meant to achieve and how they should be evaluated. On the other end of the spectrum of hiring for diversity are community colleges which, as our results show, lead the way in setting hiring standards that ensure that faculty will be able to serve all student populations. By learning from community college hiring practices, R1 institutions could become more inclusive at all levels of education, training, and research.

To address the former, one recommended practice to mitigate the biases in hiring that are inherent to any human-led evaluation is the use of systematic processes to vet applicants, which ideally would involve the use of hiring rubrics (11). Rubrics can be used to structure evaluation discussions within hiring departments and provide a more objective process for evaluating candidates. With this in mind, we hope that the ACRA rubric will be used by faculty hiring committees to help mitigate biases in hiring.

### 3. Limitations and future directions

Although the qualitative study allowed us to significantly improve the definitions of the qualifications that were required for faculty positions and of the levels of achievement necessary, there is still much to learn about hiring requirements for faculty positions.

The next step of this study will be to expand our quantitative wave to a larger sample of faculty. Expanding the sample will allow us to proceed with additional structural and external validation of the instrument, which will provide us with a sense of which qualifications co-vary among themselves and to identify potential groups of qualifications (13). A larger faculty sample will allow us to identify the predictors of hiring practices based on institutional and departmental characteristics, and to determine if there are any discipline-specific differences among faculty hiring practices. Such a study is underway and we encourage readers to collect data at their institution and within their research communities (bit.ly/facultyACRA).

It is important to note that, in the interviews, faculty were asked to reflect on the practices used by the faculty who had served on hiring committees with them, not just their own opinions. However, the current version of ACRA only represents the practices of a small number of departments in the country. In addition, it relies on the self-awareness and metacognition of faculty when it comes to the hiring process. It is possible that faculty members may not always think of all the stages of selection involved in the hiring process when asked to reflect on their hiring priorities. For example, when discussing the process for hiring participants, one R faculty mentioned fellowships as a selection criterion when sorting applicants at one of the earliest stages of application, a step that is sometimes executed by an administrative assistant: “*Fellowships is a big one to sort on.”* Interestingly, this step was originally omitted from the discussion about fundability by this faculty member. If this finding was confirmed with other R faculty members, it could suggest that prior funding could be a more significant contributing factor to hiring decisions than faculty in our sample had realized.

In addition, the ACRA does not predict the *success* of new faculty once they are hired. From our discussion with faculty, until institutions start taking a systematic approach to aligning hiring practices to success metrics for their faculty, the ACRA rubric will merely reflect what trainees must *demonstrate* to get hired, not necessarily what it takes to *succeed* as a new faculty, an important distinction to make when presenting the ACRA rubric to trainees. It also means that the ACRA rubric will need to be updated to reflect changes in hiring practices in years to come. Future studies of hiring practices using ACRA could help determine if institutional hiring practices have changed over time. It would also be interesting to determine whether the use of ACRA-like rubrics to make systematic hiring decisions results in an improvement in the retention and success of faculty hires over time.

## Acknowledgments

### General

We are appreciative of the many faculty who have contributed their time and expertise to the development of ACRA through interviews. We would like to thank Kelly Albus, Andrea Goldfien, Bill Lindstaedt, Victoria McGovern, Thi Nguyen, Naledi Saul, Allyson Spence, and Allison Hunter for their scientific and technical support in developing and conducting this project. We also thank Rachel Care, D’Anne Duncan, Michael Matrone, Michael Mullen, Gabriela Monsalve, Sapna Puri, Alexandra Schnoes, Tina Solvik, Elizabeth Watkins, Sumitra Tatapudy and the UCSF Trends, Issues and Programs meeting attendees for their support of the project and/or for feedback on our findings. A special thanks to Stephanie Gardner, Diane Ebert-May and John Vasquez for their thoughtful feedback on the manuscript, as well as the members of the Graduate Career Consortium Education Research Committee including Kelly Ahn, Amanda Bolgioni-Smith, Andrew Green, Christine Kelly, Natalie Lundsteen, Chris Smith, and Shoba Subramanian.

We would like to thank the American Association of American Colleges for awarding the ACRA rubric the first prize of the 2019 Innovations in Research and Research Education Award.

## Author contributions

L.C. designed the Prototype ACRA, designed the qualitative study with J.D. and R. M., conducted pilot interviews with J.D., designed the interview guide and Interview ACRA with J.D., conducted the first cycle of analysis of the interviews with J.D. and conducted the second cycle of analysis, designed the quantitative study, conducted the quantitative analysis, and wrote the manuscript.

J.D. designed the qualitative study with L.C. and R. M., conducted all interviews, designed the interview guide and Interview ACRA with L.C., conducted the first cycle of analysis of the interviews with L.C. and provided feedback on the paper.

R.M. designed the qualitative study with L.C. and J.D., provided guidance on the development of the interview guide and qualitative data analysis, assisted L.C. by editing of the manuscript.

