## Supplemental Files for "The Academic Career Readiness Assessment: Clarifying training expectations for future life sciences faculty"

### SUPPLEMENTARY MATERIALS

#### S1 - Sources used for the development of the Prototype ACRA rubric.

| **Sources used for developing the Prototype ACRA rubric** |
|---|
| 1. Job search books (*1–4*); Published articles referring to the hiring process for life science/biomedical faculty, as well as the tenure process and to graduate training (*5–8*) |
| 2. Faculty job descriptions posted on various websites. Seven job descriptions (three R1 job descriptions, one Master's (M) granting institution job description, three Liberal Arts College job descriptions (LAC)) were obtained from four websites (*9–12*) |
| 3. Feedback from three faculty (two at R1 institutions, one at a Liberal Arts College) on candidates' application materials: "*If you saw this application as a hiring committee member, would you consider this trainee as a worthwhile candidate? If so, why? If not, why not? Do you have any constructive feedback for the candidate?"* |
| 4. Expertise of career and professional development experts in our office with experience with hiring at community college, R1 and Primarily Undergraduate Institutions hiring practices |

#### S2- Interview faculty sample, by institution type

[1] The Basic classification of the 2015 Carnegie Classification of Institutions of Higher Education (*13*), [2] Awarded > 20 research/scholarship doctoral degrees during the year; [3] Awarded > 50 master's degrees and < than 20 doctoral degrees during the year; [4] Institutions where baccalaureate or higher degrees represent > 50 % of all degrees but with < than 50 master's degrees or 20 doctoral degrees; [5] Institutions at which the highest level degree awarded is an associate's degree; [#] Two of the R1 and two of the BAC institutions were represented by one faculty member each.

| **2015 Carnegie Basic Classification** [1] | **Group** | **Abrev.** | **Inst.** | **Faculty** |
|---|---|---|---|---|
| Doctoral Universities [2] - Highest Research Activity | R | R1 | 5[#] | 4 |
| Doctoral Universities [2] - Higher Research Activity | RT | R2 | 2 | 2 |
| Doctoral Universities [2] - Moderate Research Activity | RT | R3 | 2 | 2 |
| Master's Colleges & Universities [3] - Larger Programs | RT | M1 | 2 | 2 |
| Master's Colleges & Universities [3]- Medium Programs | RT | M2 | 1 | 1 |
| Baccalaureate Colleges [4] | RT | BAC | 3[#] | 2 |
| Associate's Colleges [5] (Community Colleges) | T | CC | 5 | 5 |
| | | **Total** | **20** | **18** |

#### S3 - Overview of Interview slides

SLIDE 1: Introductory slides, presents PI and researcher
SLIDE 2: Our focus:

- Faculty job search preparation for biomedical trainees
- Goal: Increase trainee knowledge of what skills/experiences help you get hired at different types of institutions:
  - Community Colleges
  - Liberal Arts
  - R1
  - Masters-granting

SLIDE 3: Central question for this interview: "In your experience hiring assistant professors, which of a candidate's skills or experiences contribute significantly to hiring decisions?" Will ask you to consider 16 possible factors

SLIDE 4: Which of these factors **contributes significantly** to hiring decisions?

- Teaching Experience
- Teaching Philosophy
- Scientific Communication
- Commitment to Diversity
- Collegiality
- Fit
- Mentoring
- Research with Undergrads
- Scientific Independence
- Fundability
- Recognition/Reputation
- Personal connections
- Collaboration
- Scholarship
- Scientific Vision
- Research Feasibility

SLIDE 5: What level of achievement do you hire at? [*QUALIFICATION scale is listed here]*
On the following slides, the interviewer then presents each of the scales selected by the subject in slide 3. Occasionally, some qualifications will be presented alongside one another.

FINAL SLIDES: Other important factors we've missed? Any other faculty we might talk to?

#### S4- Survey faculty sample, by institution type

[1] The Basic classification of the 2016 Carnegie Classification of Institutions of Higher Education, [2] Awarded > 20 research/scholarship doctoral degrees during the year or > 30 professional practice doctoral degrees in at least 2 programs; [3] Awarded > 50 master's degrees and < than 20 doctoral degrees during the update year; [4] Institutions where baccalaureate or higher degrees represent > 50 % of all degrees but with < than 50 master's degrees or 20 doctoral degrees; [5] Institutions at which the highest level degree awarded is an associate's degree.

| 2016 Carnegie Basic Classification [1] | Abrev. | Faculty | Group |
|---|---|---|---|
| Doctoral Universities [2] – Very high research activity | R1 | 3 | R |
| Doctoral Universities [2] - High Research Activity | R2 | 2 | RT |
| Doctoral/Professional Universities [2] | D/PU | 2 | RT |
| Master's Colleges & Universities [3] - Larger Programs | M1 | 3 | RT |
| Master's Colleges & Universities [3]- Medium Programs | M2 | 1 | RT |
| Baccalaureate Colleges [4] | BAC | 3 | RT |
| Associate's Colleges [5] (Community Colleges) | CC | 3 | T |

| | | |
|---|---|---|
| | **Total** | **17** |

**S5- Inter-rater reliability (IRR) using the Final ACRA rubric for T, RT and R institutions ("Group").**
Institution subtype refers to the 2016 Carnegie Classification for the institutions to which belonged the two faculty members in each pair. IRR refers to the percentage agreement between raters on whether Final ACRA qualifications are required at their institution.

| Group | Institution subtype | IRR | Divergent qualifications |
|---|---|---|---|
| T | Associate's colleges (Community Colleges) | 86.7% | Collaborations, Recommendations, |
| RT | Baccalaureate colleges | 80% | Collegiality, Funding Plan, Teaching Experience |
| RT | Master's Colleges & Universities - Larger Programs | 100% | N/A |
| R | Doctoral Universities – Very high research activity | 93.3% | Teaching Practices |

**S6 - The hypothesized qualifications required to receive a faculty job offer at different types of institutions are represented in the Prototype Academic Career Readiness Assessment rubric (Prototype ACRA).**
Symbols indicate the hypothesized hiring levels of one of three categories of U.S. institution (Circles: research-intensive institutions; Squares: research and teaching-focused institutions; Triangles: teaching-only institutions). The rubric presented ten qualifications, each featuring four levels of achievement, from basic (lowest level required by an institution to receive an offer for a faculty position) to advanced (highest level required).

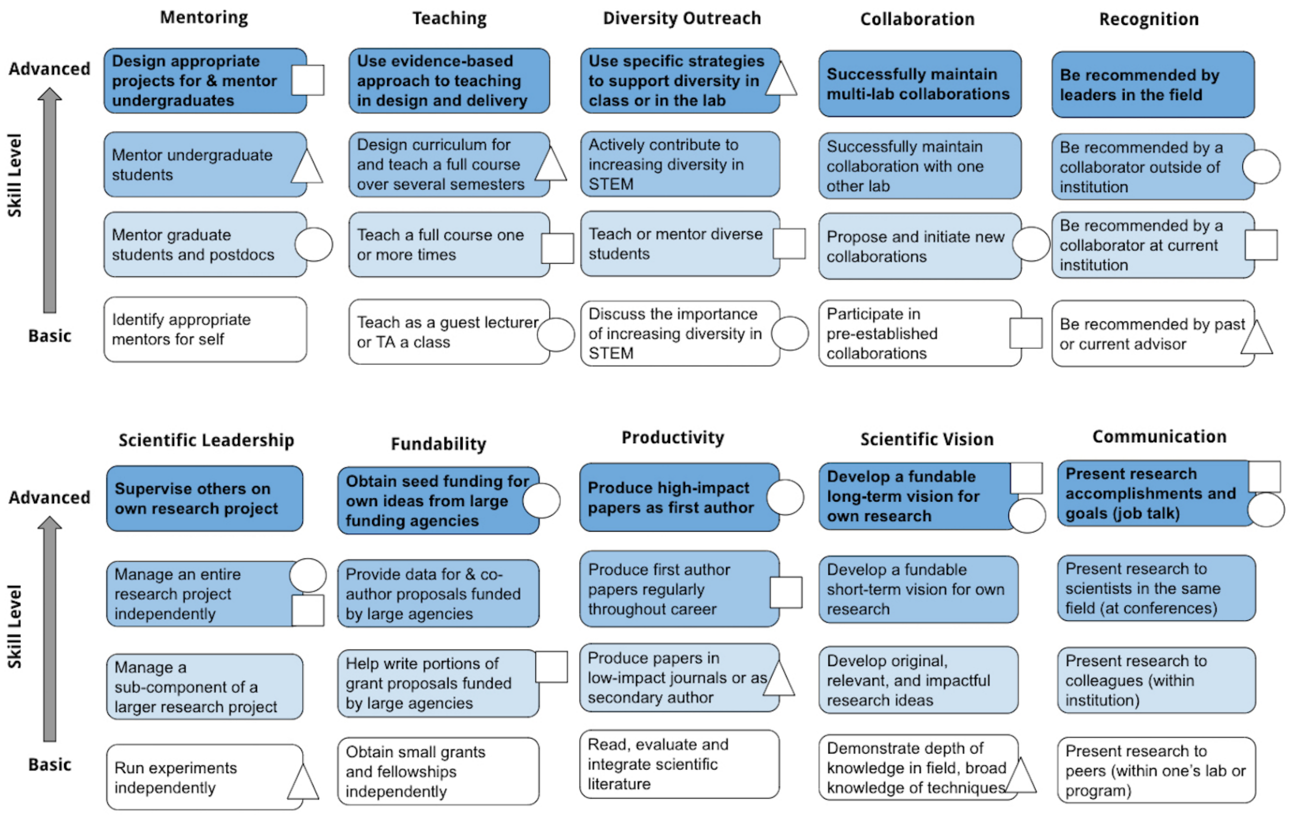

**S7 - Definitions of qualifications and corresponding sources used in the development of the Prototype ACRA rubric.** Citations (in parenthesis) refer to publications and the corresponding quote or section in said article. "Faculty" refers to feedback received from faculty on application materials: two R1 (A and B) and one Liberal Arts College (LAC) faculty members. "Job description" refers to seven sample job descriptions, collected in 2014 during the development of the Prototype ACRA rubric: three R1 job descriptions, one Master's (M) granting institution job description, three Liberal Arts College job descriptions (LAC). For more details about the sources mentioned here, refer to Supplementary Materials S2 and S3.

| **Qualifications** | **Sources** |
|---|---|
| **Vision**<br>*Develop broad and depth in scientific knowledge, develop a research vision* | (5): "*Broad Knowledge of Discipline, Broad Interdisciplinary Knowledge of Experimental Approaches, Depth of Knowledge in Your Field, Ability to Have New Insights: Being Creative, Ability to Choose Important Research Problems*"<br>(8) "*Give an overview of your research agenda, including short- and long-term objectives. (...) Be sure to convey to your audience why the work is important and why you can make a difference in the field*"<br>**R1 faculty A:** Identify the gap in the field the candidate would be filling, a unique area to which they would contribute.<br>**LAC job descriptions 1 & 2**: Candidate will develop an active research program that can involve undergraduate students.<br>**R1 job description 1:** Submit "*a 1-2 page summary of research accomplishments, a 1-2 page perspective on future research plans*"<br>**R1 job description 2:** "*Candidates will be expected to develop an exceptional research* |

| | |
|---|---|
| | *program*"<br>**R1 job description 3:** "*We seek creative scientists using cutting edge technologies and experimental systems"* |
| **Productivity**<br>*Regular publications, with impact and authorship* | (5): "*Ability to Read, Evaluate and Integrate Scientific Literature*"<br>(8): "*Components of a job application: Publications (...) Statement of your research accomplishments"*<br>**R1 faculty B:** Best to apply when all postdoctoral publications are out.<br>**R1 job description 3**: "*Applicants should submit (...) a list of publications*" |
| **Fundability**<br>*Demonstrate ability to write and obtain funding* | (8): "*Components of a job application: Major sources of independent funding, (...) Awards and honors, including pre- and post-doctoral fellowships"*<br>**LAC faculty**: Funding is a good thing to mention. Transitional awards should be mentioned early on in the materials.<br>**R1 job description 2:** "*Candidates will be expected to develop and maintain extramural funding for this research program*"<br>**R1 job description 3:** "*Successful candidates will be expected to establish an independent research program*" |
| **Teaching**<br>*Teaching experience, curriculum design experience, use of evidence-based practices* | (6): "*Elements of an outstanding teaching demonstration (...) the use of active learning*"<br>(7): "*Offers evidence of practice*"<br>(8): "*Components of a job application: Teaching Experience*"<br>**LAC faculty**: Lead with teaching experience.<br>**LAC job description 2**: Submit evidence of demonstrated or potential excellence in undergraduate instruction such as complete sets of teaching evaluations.<br>**R1 job description 2:** "*Candidates will be expected to teach undergraduate and graduate courses and develop new courses*"<br>**R1 job description 3:** "*Successful candidates will be expected to (...) contribute to the educational mission of the Department and the School."*<br>**M job description:** "Teach undergraduate students in a variety of both introductory and upper level courses. Work with faculty to strengthen the existing degree programs" |
| **Mentoring**<br>*Have experience mentoring students, and potentially demonstrate ability to design projects for students* | (8): "*Your role as a laboratory leader*"<br>**LAC faculty:** identify undergraduate students in publication list.<br>**LAC job description 1**: Include a statement of research plans that includes how undergraduates might be included.<br>**R1 job description 2:** "*Candidates will be expected to advise graduate students and post-graduate researchers*"<br>**LAC Job description 3:** "*Candidates who are firmly committed to undergraduate education, including the involvement of undergraduates in productive research programs of nationally recognized quality."* |
| **Communication**<br>*Present research in different contexts* | (5): "*Ability to Think Clearly: Effective Oral and Written Communication"*<br>(8): "*Components of a job application: Invited keynotes and applications"* |
| **Recognition**<br>*Obtain recommendations from mentors and leaders in the field* | (8): "*Components of a job application: References, (...) Letters of Recommendations"*<br>**M job description**: Submit *"The names and contact information for three current or former supervisors or colleagues who can serve as references with respect to your academic experience and Successes."*<br>**LAC job description 2:** Submit "*three letters of reference"* |
| **Leadership**<br>*Manage research* | (5): "*Ability to Plan and Execute Experiments*"<br>(8): "*Build and manage teams*" |

| | |
|---|---|
| *projects, and potentially supervise research teams* | **R1 job description 2:** "*Candidates will be expected to advise graduate students and post-graduate researchers*" |
| **Collaboration**<br>*Initiate and manage collaborations* | (*8*): "*Setting up a collaboration*"<br>**LAC faculty:** Collaborations are good things to mention.<br>**R1 faculty B:** Propose collaborations with the faculty at the target institution. |

**S8 - Resources used to develop the four levels of achievement for the additional qualification in the Prototype ACRA**

| | |
|---|---|
| **Diversity Outreach**<br>*Engage with diverse student populations, potentially using effective practices* | (*7*) *"Is attuned to differences in student ability, learning styles, or level"*<br>**LAC faculty:** Describe sensitivity to needs of URM students.<br>**LAC job description 1**: "*We are particularly interested in applicants who will strengthen the departmental commitment to students from underrepresented groups, and candidates committed to teaching a diverse student body.*"<br>**LAC job description 2;** applicants should explain how their pedagogy will serve to create and sustain an inclusive learning environment.<br>**LAC job description 3:** "*especially interested in applicants who can contribute to the diversity of the College and its excellence through their research, teaching and/or service.*" |

#### S9 - Interview ACRA rubric

| | Level 1 | Level 2 | Level 3 | Level 4 |
|---|---|---|---|---|
| **Teaching Experience** | No teaching experience | Has given a guest lecture or served as a teaching assistant | Has taught a full course one or more times | Has designed and taught a course |
| **Teaching Philosophy** | Shows intent to serve department's teaching needs | Reflective about effectiveness of approaches used for student population | Uses validated approaches grounded in the literature | Collects data on student learning and uses an iterative process to improve curriculum |
| **Scientific Communication** | Can present research clearly to labmates | Can present research clearly to colleagues at your institution who are less familiar with research area. | Can present research clearly to scientists at conferences | Can deliver an effective talk to an educated non-expert audience (i.e., in a job talk) |
| **Commitment to Diversity** | Can explain diversity issues in STEM fields | Has taught or mentored diverse students | Actively contributes to diversity efforts | Uses strategies to support diversity in class or in lab |
| **Fit** | Shows an understanding of institution's needs (student population, mission, etc.) | Shows willingness to meet institution's needs | Demonstrates that experience is a good fit with institution's needs | Demonstrates strong potential forgetting tenure |
| **Collegiality** | Does not ask any questions about other faculty's work. | Shows interest in new ideas and other faculty's work. | Shows potential for interacting well with faculty (fits in with departmental culture) | Shows potential for developing successful collaborations with other faculty at the institution |
| **Scholarship** | Can read, evaluate and integrate scientific literature | Has produced low-impact, 2nd author papers | Has produced 1st author papers regularly | Has produced high impact, 1st author paper |

| | Level 1 | Level 2 | Level 3 | Level 4 |
|---|---|---|---|---|
| **Research with Undergrads** | Has published data collected with the research system (but not yet collected with undergrads) | Has collected preliminary data with undergraduate mentees | Has presented posters with undergraduate mentees | Has published data with undergraduate mentees |
| **Mentoring** | Expresses enthusiasm for mentoring students on how to do research | Can articulate their approach to mentoring | Has some experience mentoring students or trainees | Has extensive experience mentoring students or trainees |
| **Scientific Independence** | Has run experiments independently | Has managed a sub-component of a larger project | Has managed an entire research project independently | Has managed a research team (people, projects, & administrative tasks) |
| **Fundability** | Has obtained small grants or fellowships | Has written portions of proposals funded by large agencies | Has co-authored proposals funded by large agencies | Has obtained funding for own ideas from large agencies |
| **Recognition** | Recommended by past or current advisor | Recommended by collaborator at current institution | Recommended by external collaborator | Recommended by leaders in the field |
| **Personal connections** | Search committee member knows the candidate's **advisor** | Search committee member has heard candidate's **advisor** talk about the candidate's research | Search committee member has heard the **candidate** speak at a conference | Search committee member knows the **candidate** personally |

| | Level 1 | Level 2 | Level 3 | Level 4 |
|---|---|---|---|---|
| **Collaboration** | Has **participated in** pre-existing collaborations | Has **led** collaborative projects | Has **initiated & led** collaborative projects | Has **ongoing collaborations** that could be taken to the new institution |
| **Scientific Vision** | Demonstrates knowledge of field | Has developed original, impactful research ideas | Has developed a short-term vision for own research | Has developed a long-term vision for own research |
| **Research Feasibility** | Research program is not tailored to the resource constraints of the institution nor to the student population | Research program is tailored to resource constraints of the institution | Research program is tailored to resource constraints *and* to student population | Level 3 + Some aspects of the research have already been done with relevant student population |

#### S10 - ACRA rubric footnotes

1. An additional level was suggested as being advantageous, but not required for a faculty position at T and RT institutions: "Level 4 & Candidate has collected evidence on student learning (discipline-based education research)."
2. Including curricular and management responsibilities and substantial interactions with undergraduate students. These candidates should be able to demonstrate their effectiveness through course evaluations, their philosophy through their teaching

statement and their potential in the teaching demonstration. These candidates should also show potential for being mentored as a new faculty.

3. Including curricular responsibilities (syllabus, lecture, assignment and exam development). Candidates should be able to demonstrate their classroom management skills in the interview.
4. In particular, lower-division undergraduate students.
5. For e.g., at an RT institution as a Visiting Assistant Professor, and at a T institution as an Adjunct Faculty.
6. An additional level was suggested as being advantageous, but not required for a faculty position at T institutions: “Level 4 & Candidate has collected classroom or institutional data around equity and has engaged in efforts to create an equitable learning environment for students.”
7. The research program should include projects that are compatible with the institution’s typical course schedule, diverse levels of research skills (novice vs. advanced) and education levels (freshman, senior, Master’s student).
8. At R1 and R2 institutions, this is mainly applicable to candidates who come from large, highly funded labs (for e.g., HHMI-funded labs) and whose program scope needs to be tailored to the resources available to a junior PI in the first few years, as they grow their team.
9. At R1 and R2 institutions, this is mainly applicable to candidates who require high-end equipment.
10. At some R1 and R2s, this may mean the absence of a certain type of facility, or a lack of space in the animal facility, or the distance from the medical school for work on human subjects or samples.
11. At RT institutions, where start-up funds are limited and core facilities often nonexistent, research requiring some animal models or expensive equipment may not be feasible. Candidates are expected to tailor their research plan the specific resources of each institution.
12. The word “and” here refers to the frequency of publications during a candidate’s training. “And” indicates that the hiring committee is looking for a consistent pace of publication, both during graduate school and postdoctoral training. Some RT institutions indicated that they were looking for candidate has produced at least one first author paper during postdoc or PhD (regardless of impact or frequency)
13. The number of papers required to get a faculty job offer was related to the level of research at that institution per the Carnegie Classification, i.e. R2 institutions required a dozen publications, if not of high impact, while R3 and M1 institutions required “a couple” of publications (for e.g. two first author publications during the PhD and two during the Postdoc).
14. Faculty have reported that hiring committees often discuss a paper’s contribution to the field beyond the impact factor of the journal in which it is published, considering important journals to specific subfields, and work that shows potential to advance science, as well as the creativity of the research and the novelty of the findings.
15. The faculty members in our sample did not necessarily require these types of publications, but did describe a tension within hiring committees with other faculty members around this. Some suggested that there may be an implicit bias in favor of candidates with these types of publications.
16. For e.g., research question is exciting, or methodology is cutting edge. The emphasis at R1, R2 and RT institutions is on getting other faculty excited about this research. In addition, at RT institutions, faculty members will be looking for a research program that is exciting for students.
17. This can include having previously identified a gap in the field and developed and conducted experiments to fill this gap as a postdoctoral scholar, or having previously collected preliminary data to demonstrate the technical feasibility of the program.
18. R3, M1 and M2 institutions prefer a candidate that has at least given some thought to the type of funding program that could support their research plan.
19. At R1 and R2 institutions, the research program is assessed through the lens of an R01 grant study section. Candidates are expected to demonstrate creativity, as well as to discuss the potential impact of their research program on their field.
20. Candidates are expected to present specific aims that are within the scope of R01-funded grants.
21. This can include specific aims for large grants other than the first R01 grant.
22. In the case where the program relies on collaborations, these collaborations will be maintained in new position.
23. This stage corresponds to a shift from the postdoctoral to the faculty identity. In addition to having a clear research vision and strategy, the candidate will need to demonstrate an ability to envision alternative approaches, evaluating results, and setting new directions for a project.
24. Because the projects are distinct, or because the advisor and candidate plan to maintain clear boundaries.
25. This enthusiasm is more impactful when expressed by a scientist who is not typically as enthusiastic about applicants, and when it is personalized, i.e. specifically describes the candidate, their accomplishments and their potential. In addition, having the recommender reach out directly to the search committee can be influential. Note that some RT (but no R) institutions have reported following up with candidates who are missing a recommendation letter from one of their PIs.
26. Either through personal connections or because the PI has a strong reputation in the field.
27. At R institutions, this involves demonstrating curiosity for other faculty’s work and ideas, while at RT institutions, it involves getting along with colleagues. At T institutions, this fit is often demonstrated through other competencies, like Teaching Potential, Teaching Experience or Commitment to Serving Diverse Students, as a sort of compound competency.
28. At RT institutions, this may involve sharing of equipment, space and materials. At T institutions, it involves discussing and sharing curriculum and course materials, and discussing interaction with students.
29. At RT institutions, this may be collaborating on research or educational projects. At T institutions, it may look like team-teaching, collaborating with a colleague on the development of new curriculum or the development of a learning community.

30. For example, when applying for position at a T institution, the candidate has sought out opportunities to teach to align with the institution's teaching mission.
31. At R1 and some R2 institutions, this means that the candidate's research expertise does not compete with existing research programs. At RT, T and some R2 institutions, the teaching discipline potentially covered by the candidate fill a gap in the department.
32. At RT institutions, this means a high teaching load. At R2 institutions, it means a high teaching and research load.

9. ChronicleVitae. *ChronicleVitae for higher ed jobs, career tools and advice*, (available at https://chroniclevitae.com/job_search/new).

10. Cell Biology Jobs - American Society for Cell Biology, (available at https://jobs.ascb.org/).

11. Academic Jobs Online, (available at https://academicjobsonline.org/ajo).

12. Welcome to the Carleton College Applicant Portal Site | Home, (available at https://jobs.carleton.edu/).

13. Carnegie Classifications, (available at http://carnegieclassifications.iu.edu/index.php).
